# Novel Asgard archaea phylum Hermodarchaeota degrade alkanes and aromatics via alkyl/benzyl-succinate synthase and benzoyl-CoA pathway

**DOI:** 10.1101/2020.10.19.346239

**Authors:** Jia-Wei Zhang, Hong-Po Dong, Li-Jun Hou, Yang Liu, Ya-Fei Ou, Yan-Ling Zheng, Ping Han, Xia Liang, Guo-Yu Yin, Dian-Ming Wu, Min Liu, Meng Li

## Abstract

Asgard superphylum is composed of a group of uncultivated archaea that are deemed the closest relatives of eukaryotes. These archaea are widely distributed in anaerobic environments and suggested to be important players in carbon cycling of sediments. Alkanes and aromatics are refractory organic compounds and abundant in sediments. However, little is known about degradation of these compounds by Asgard archaea to date. Here, we describe a previously unrecognized archaeal phylum, Hermodarchaeota, affiliated with the Asgard superphylum. The genomes of these archaea were recovered in metagenomes from mangrove sediments, and were found to encode alkyl/benzyl-succinate synthases and their activating enzymes that are similar to those found in alkanes-degrading sulfate-reducing bacteria. Hermodarchaeota also encode enzymes for alkyl-coenzyme A and benzoyl-coenzyme A oxidation, and the Wood–Ljungdahl pathway, as well as nitrate reductases. Furthermore, transcripts for these enzymes have been frequently detected in metatranscriptomes from mangrove sediments. This indicates that members of this phylum are able to anaerobically oxidize alkanes and aromatic compounds, coupling the reduction of nitrate. Genes encoding 16S rRNA and alkyl/benzyl-succinate synthases analogous to those in Hermodarchaeota were identified in a range of marine and freshwater sediments. These findings suggest that Asgard archaea capable of degrading alkanes and aromatics via formation of alkyl/benzyl-substituted succinates are ubiquitous in sediments.

---

Alkanes and aromatic hydrocarbons are abundant and prevalent in the environment. Although they are major components of petroleum, living organisms are also their important sources. These compounds are inactive in molecular structure, which makes them relatively inert substrates. Under aerobic conditions, alkanes can be oxidized by monooxygenase or dioxygenase in aerobic microorganisms, which use oxygen to supply a reactive oxygen species. The resulting alcohols are further oxidized to aldehydes by dehydrogenases, which are then converted to fatty acids [1]. However, the anaerobic oxidation of alkanes requires further study. Some sulfate-reducing bacteria (SRB) are capable of utilizing n-hexadecane [2], propane, and n-butane [3]. SRB activate alkanes through the addition to fumarate, producing alkyl-substituted succinates [3]. This mechanism is analogous to the anaerobic activation of aromatic hydrocarbons, yielding benzylsuccinate as the first intermediate, in the denitrifier *Thauera aromatica* [4]. For Archaea, only a few anaerobic species are currently known to have the ability to grow on hydrocarbons, with the exception of methane. *Ferroglobus placidus* can degrade benzene at 85°C, coupling reduction of Fe(III) [5]. The thermophilic sulfate-reducing archaeon, *Archaeoglubus fulgidus*, was shown to oxidize long-chain n-alkanes anaerobically. It was inferred that *Archaeoglubus fulgidus* might activate alkane through binding to fumarate, which is catalyzed by alkylsuccinate synthase [6]. Recently, a thermophilic archaea (*Candidatus Syntrophoarchaeum*) was shown to activate butane via alkyl-coenzyme M formation under anaerobic conditions, which is similar to anaerobic activation of methane. Genes encoding a similar methyl-coenzyme M reductase (MCR) complex have also been identified in genomes of uncultivated Bathyarchaeota, Hadesarchaeota [7] and Helarchaetota [8].

The recently discovered Asgard superphylum is a group of uncultivated archaea with many eukaryotic features including six distinct phyla: Loki-, Thor-, Odin-, Heimda-, Hel-, and Gerd-archaeota [8–10]. These archaea possess so-called eukaryotic signature proteins (ESP), which, in eukaryotes, are involved in membrane-trafficking processes, vesicle biogenesis and trafficking, cytoskeleton formation and remodeling, endosomal sorting complexes required for transport-mediated protein degradation, and endosomal sorting; therefore, they are deemed representative of the closest archaeal relatives of eukaryotes [9]. Diversity investigations have revealed that Asgard archaea are widely distributed in various anoxic environments, including mangrove sediments, estuarine sediments, freshwater sediments, hydrothermal habitats, marine sediments, cold seeps, hot springs, mud volcanos, and soils [8, 9, 11, 12]. Genomic analysis suggests that Asgard archaea may primarily be organoheterotrophs but some of them, such as Lokiarchaeota, Thorarchaeota and Gerdarchaeota, may also be mixotrophs, which can perform carbon fixation via the Wood–Ljungdahl pathway (WLP) [9, 12]. The versatile lifestyles of Lokiarchaeota, Thorarchaeota and Gerdarchaeota have been supported by metatranscriptomics [12]. More recently, Helarchaeota from hydrothermal deep-sea sediments is suggested to have the potential to oxidize short-chain hydrocarbon using MCR-like enzymes [8], similar to the butane-degrading archaea *C. Syntrophoarchaeum*. These results underscore the roles of Asgard archaea in global carbon cycling.

Here, we present the discovery of metagenome-assembled genomes recovered from anoxic mangrove sediment belonging to a new Asgard phylum that has the potential to carry out anaerobic oxidation of alkanes and aromatic compounds through binding to fumarate producing alkyl/benzyl-substituted succinates, which further extends our knowledge on carbon metabolism of Asgard archaea.

## Methods

### Sample collection and metagenomic sequencing

Six sediment samples were taken from the site H0 at mangrove swamps on Techeng Island, Zhanjiang, Guangdong, China on November 25, 2018 **(Supplementary Fig. 1)**. The site was located in a 500-year-old national nature reserve for mangroves. Two cores with 1 m depth (H02 and H03) were collected using a peat sampler and they were approximately 1 m apart. The freshly cores were divided into surface (15–20 cm), intermediate (40–45 cm), and bottom (95–100 cm) sections for further analysis, and stored at −80°C. Detailed data of these sediment samples is provided in **Supplementary Table 1**.

Genomic DNA was obtained from about 10-15 g of sediment samples with PowerSoil DNA Isolation Kit (MoBio Laboratories, Carlsbad, CA, USA). Metagenomic sequence data for the six sediment samples was produced using Illumina HiSeq 2500 instruments at Guangdong MagiGene Technology Corporation (Guangzhou, China). The amount of raw sequence data was approximately 60 Gbp for each sample (2 × 150 bp).

### Metagenome assembly and genome reconstruction

Raw reads were dereplicated and trimmed to get rid of adaptors and low-quality reads with Trimmomatic [13]. The high-quality metagenomic sequences were assembled using either MEGAHIT [14] with the following parameters: --k-min 27--k-max 127--k-step 10, or IDBA-UD [15] using the parameters: -mink 55-max-k105-steps 10. Genome binning was performed using an association of MetaBAT [16], Emergent Self-Organizing map (ESOM) [17], MaxBin [18], and CONCOCT [19]. Briefly, contigs with length < 1,500 bp in each assembly were removed and the coverage information of the remaining contigs was computed with Bowtie2 [20]. Genome binning was then performed based on the coverage information and tetranucleotide frequency by MetBAT [16], MaxBin [18], and CONCOCT [19]. Contamination in bins was checked carefully and removed using ESOM [17] and mmgenome toolbox (http://madsalbertsen.github.io/mmgenome/). For ESOM [17], these bins were sheared into short fragments (5 to 10 kb) and grouped according to the tetranucleotide frequency in ESOM [17] (**Supplementary Fig. 2**). Subsequently, the resulting groups were sorted manually and each group represented one single bin. These bins were recovered with getClassFasta.pl. Two control genomes were added to raise ratio of signal to noise and improve binning (**Supplementary Fig. 2**); complete recovery was possible. The GC content of contigs in each bin was computed using mmgenome R package (https://github.com/MadsAlbertsen/multi-metagenome/blob/master/R.data.generation/calc.gc.pl). The GC content and coverage information for contigs of the bins were used to draw a scatter plot in which the clustered spots representing one single bin were detected with the mmgenome R package. Finally, the bins obtained were curated manually to remove contamination based on multi-copy marker genes. CheckM [21] was used to assess completeness and contamination of these genome bins. As a result, seven high-quality genome bins generated using the MEGAHIT [14] assembly, belonging to a unknown phylum of Asgard archaea, were sorted for further analysis (**Supplementary Table 2**). These bins were named Hermodarchaeota. The AAI calculated by CompareM (https://github.com/dparks1134/CompareM) was applied to compare the discrepancies between Hermodarchaeota bins and known Asgard genomes.

### 16S rRNA gene analysis

Among above-mentioned seven Hermodarchaeota bins, only h02m_131 contained one 16S rRNA gene sequence (506 bp in length). To find longer 16S rRNA genes, raw reads were mapped to all Asgard bins using Bowtie2 [20] with default parameters. The resulting reads were combined with unbinned reads. Subsequently, these reads were assembled using metaSPAdes [22] with the parameters: -k 55, 65, 75, 85, 95. The scaffolds produced were binned using procedures as described above. After contamination was removed, we recovered one bin containing a 16S rRNA gene with a length of 1066 bp (h02s_26) (**Supplementary Table 3**). This bin and h02s_124 is highly similar each other (98.86% AAI), likely representing the same species. In order to search for homologues of Hermodarchaeota 16S rRNA gene sequence, raw reads were mapped to the Silva database [23] and then assembled using MATAM [24] to generate a database containng all 16S rRNA gene sequences in the samples. The 16S rRNA gene sequence of h02s_26 was used to identify homologs within the database using Blastn [25]. Two additional 16S rRNA gene sequences were detected, which had > 95% identity with the query sequence (**Supplementary Table 3**). Furthermore, comparison to the existing 16S rRNA genes of Asgard archaea was performed with Blastn [25].

16S rRNA genes of Asgard including Hermodarchaeota were aligned along with those of Euryarchaeota with MAFFT-L-INS-I [26] and pruned using trimAl [27]. For 16S rRNA genes with genome representatives, genome information of their taxa is shown in **Supplementary Table 4**. These Asgard 16S rRNA genes without genomic information are derived from a previous study [10]. Maximum-likelihood (ML) tree was conducted with IQtree [28] (v.1.6.12) using the GTR+F+I+G4 model with 1,000 ultrafast bootstraps.

### Phylogenomic analysis of concatenated ribosomal proteins

A suit of representative Archaea were used in the analysis; they were carefully selected according to a taxa list reported previously [9], consisting of 72 archaea. The homologs of 56 ribosomal proteins in eight Hermodarchaeota bins and representative archaeal reference taxa were identified through Archaeal Cluster of Orthologous Groups (arCOGs) using Blastp [29] (cutoff: e-value, 1e-5). The full lists of organisms and ribosomal proteins used for phylogenomic inference are shown in **Supplementary Table 4 and Supplementary Table 5**, respectively. Each of the 56 ribosomal proteins was aligned using MAFFT-L-INS-I [26], checked manually, and pruned using trimAl [27] under automated 1 or BMGE [30] with BLOSUM30 matrix. Trimmed alignments were concatenated, and then further aligned with MAFFT-L-INS-I [26] and pruned using BMGE [30] or trimAl [27] to generate a final alignment. An ML phylogeny was inferred with IQtree [28] using the LG+C60+F+G+PMSF model and ultrafast bootstrapping (1,000 replicates). Bayesian phylogeny was inferred using PhyloBayes [31], using the LG+CAT model. Two Markov chains were run and each chain generated 12117 trees. The maximum difference between the two chains was acquired after removing the first 100 trees (maxdiff < 0.25).

### Annotation of genome bins and metabolic reconstruction

Genes for all Hermodarchaeota bins were analyzed with Prodigal [32] (v.2.6.3) using the setting: -p meta. Prokka [33] was also used for performing gene prediction but fewer genes were obtained by Prokka [33] than Prodigal [32]. Gene annotation was done by searching against the non-redundant (nr) database (downloaded from the National Center for Biotechnology Information (NCBI) in October 2019), existing arCOGs [34], and the UniProtKB and Swiss-Prot database [35] using Blastp [29] (cutoff: e-value, 1e-5). In addition, functional annotation for coding sequences was also conducted using eggNOG-mapper [36] against the eggNOG database. To reconstruct the metabolic pathways, annotation of coding sequences was performed on the BlastKOALA server [37]. Protein conserved domains were analyzed using InterProScan [38] with default parameters. Carbohydrate-active enzyme annotation was conducted on the dbCAN2 meta server [39]. MEROPS database [40] was used to analyze peptidases of Hermodarchaeota genomes.

To identify genes related to hydrocarbon utilization in Hermodarchaeota bins, a protein database was constructed comprising enzymes involved in anaerobic hydrocarbon oxidation in the nr database. These enzymes included the α-subunit of alkylsuccinate synthase (AssA), α-subunit of the benzylsuccinate synthase (BssA), pyruvate formate lyase (pfl), pyruvate formate lyase-activating enzyme, glycyl radical enzyme, (1-methylalkyl) succinate synthase (Mas), and α-subunit of naphtylmethylsuccinate synthase (Nms). Hermodarchaeota proteomes were queried against this database using Blastp [29] (cutoff: e-value, 1e-5). Positive hits were compared with the annotations from UniProt [35], Swiss-Prot, and Interpro [41]. Motifs or conserved residues for AssA/BssA and Ass/Bss activating proteins were analyzed by comparing these hits with known reference proteins following a previous method [6]. The bzd gene cluster in the contig with benzoyl-CoA reductase (Bcr) was analyzed according to an anaerobic benzoate degradation pathway reported previously [42].

Nitrate reductases from the nr database were used for the construction of a protein database. The predicted genes in Hermodarchaeota bins were applied to search the database by using Blastp [29] (cutoff: e-value, 1e-5). The hits with a bit score > 200 were further identified using a hmmscan search against nitrate reductases from TIGRfam [43] and Pfam [44]. The conserved residues and motifs in nitrate reductases were analyzed following a previous method [45]. These hits with the molybdopterin oxidoreductase domain and CXXCXXXC motif were regarded as likely periplasmic nitrate reductases (Nap).

Hydrogenases were identified by searching against a protein database consisting of hydrogenase sequences from nr database using Blastp [29] (cutoff: e-value, 1e-5). Hits were further verified using the HydDB [46], Interpro [41], and KEGG databases [47]. Group 4 NiFe hydrogenases and energy-converting hydrogenases related complexes were identified according to a previous reported method [48]. The CxxC motifs of hydrogenase that link H2 metal centres [49] were analyzed.

For identification of ESPs, accession number from the InterPro [41] and arCOGs provided in a ESP table reported by Zaremba-Niedzwiedzka [9] was used for searching for Hermodarchaeota proteomes annotated by InterPro [41], arCOG [50], and nr databases. The ESPs that were not matched were identified by searching all predicted genes in Hermodarchaeota bins against Asgard homologs using hmmscan.

### Metatranscriptomic analysis

Seven metatranscriptomic datasets from mangrove sediments were obtained from the NCBI database (**Supplementary Tables 6-8**). The sampling locations for these sediments are shown in **Supplementary Fig. 1**. Transcripts were identified by comparing against a database of genes involved in important metabolic processes from the Hermodarchaeota genomes using Blastn (cutoff: e-value, 1e-10). If a read was mapped to different genes, the best hit was used to determine the number of transcripts assigned to a gene [51]. The gene expression level was estimated based on the number of reads matching the gene. For the three datasets with more reads mapped to *Ass/Bss* genes, gene expression was also normalized to reads per kilobase of transcript per million mapped reads (RPKM) [52] in order for the expression levels between genes to be compared (RPKM= number of reads matching a gene × 10^9^/(gene length × total number of reads mapped to genome).

### Phylogenetic analysis of key functional genes

#### AssA/BssA sequence phylogeny

AssA/BssA sequences from Hermodarchaeota were used as seeds to search against protein sequences of Archaea and Bacteria in the nr database using Blastp [29]. In the BLAST results, sequences with an abnormal size were discarded while protein sequences that have a size ranging from 500 to 1,000 amino acids were retained. For these remaining sequences, the top 1,000 homologs with an E-value < 1e-132 and total score > 415 were selected to construct a dataset. Furthermore, these sequences in this dataset were clustered using CD-HIT [53] (≥ 90% identity), and the repeated sequences with > 90% amino acid identity were removed. Finally, 281 homologs were retrieved from Bacteria and 18 homologs from Archaea. These sequences, along with AssA/BssA sequences from Hermodarchaeota and other Asgard lineages, were arrayed with MAFFT-L-INS-I [26], and then pruned using BMGE [30] (BLOSUM30 option). An ML phylogenic tree was computed with IQtree [28] (v.1.6.12) using the LG+G4 model and ultrafast bootstrapping (1,000 replicates).

#### Nitrate reductase sequence phylogeny

Nitrate reductases of prokaryotes are classified into two major clusters, the membrane-associated prokaryotic nitrate reductase (Nar) cluster, and the periplasmic nitrate reductase (Nap)/prokaryotic assimilatory nitrate reductase (Nas) cluster. To determine evolution of nitrate reductases from Hermodarchaeota, a set of homologues of Nar, Nap, and Nas from representative prokaryotic organisms were used for the reconstruction of a phylogenetic tree according to a previous study [45]. These sequences were arrayed with MAFFT-L-INS-I [26] and pruned using BMGE [30] (BLOSUM30 option). An ML phylogeny was inferred with IQtree [28] using the best parameters obtained by internal prediction model (LG+I+G4), and ultrafast bootstrapping (1,000 replicates).

#### Bcr sequence phylogeny

Bcr sequences from Hermodarchaeota were searched against the nr database using Blastp [29]. In the BLAST results, homologs with an E-value < 2.86 e-36 were sorted. These sequences were filtered using CD-HIT [53] (≥ 90% identity) to reduce the amount of sequences. In the resulting sequences, only sequences annotated as Bcr were used for constructing the phylogeny. These sequences were arrayed using MAFFT-L-INS-I [26] and pruned using BMGE [30] (BLOSUM30 option). An ML phylogenic tree was inferred with IQtree [28] using the best parameters obtained by internal prediction model (LG +G4), and ultrafast bootstrapping (1,000 replicates).

#### Reductive dehalogenase sequence phylogeny

Functionally characterized reductive dehalogenases from bacteria have been reported previously [54]. The sequences, together with sequences of reductive dehalogenase from Hermodarchaeota and other Asgard lineages, were arrayed using MAFFT-L-INS-I [26] and pruned using BMGE [30] (BLOSUM30 option). An ML phylogenic tree was inferred with IQtree [28] using the best parameters obtained by internal prediction model (LG+I+G4), and ultrafast bootstrapping (1,000 replicates).

#### [NiFe]-hydrogenase (large subunit) sequence phylogeny

Group 3 and group 4 [NiFe]-hydrogenase from Hermodarchaeota identified by HydDB [46] was filtered by length and motifs, and only sequences with > 300 amino acids and CxxC motifs were used. Reference hydrogenases were downloaded from HydDB [46] and repeated sequences were removed by cd-hit. Sequences of hydrogenase large subunit from other Asgard archaea were derived from a previous study [55]. All hydrogenase sequences were arrayed with MAFFT-L-INS-I [26] and pruned using BMGE [30] (BLOSUM30 option). An ML phylogenic tree was performed with IQtree [28] using the predicted best model (LG+R10 for group 3; LG+R7 for group 4), and ultrafast bootstrapping (1,000 replicates).

### Environmental distribution of Hermodarchaeota Ass/Bss and 16S rRNA genes

More than 1,000 publicly available environmental metagenomes within the SRA were downloaded from the NCBI website. Reads within these metagenomes were identified by searching against AssA sequences of Hermodarchaeota using the BLASTX of DIAMOND [56] (v.0.9.25) (cutoffs: e-value < 1e-5, bit score > 50). The Hermodarchaeota Ass/Bss homologues identified are shown in **Supplementary Table 9**.

For identification of 16S rRNA gene fragments belonging to the Hermodarchaeota, reads in metagenomes from the SRA database were mapped to the 16S rRNA sequence of the h02s_26 genome using BWA [57] with default parameters. CoverM (v.0.3.1) (https://github.com/wwood/CoverM) was applied to filter reads. These reads were regarded as matched appropriately if the alignment length was ≥ 50% and the identity was ≥ 90% or 95%.

## Results and discussion

### Identification of Hermodarchaeota genomes from mangrove swamps

A total of 360 gigabases of raw sequence data from six sediment samples were obtained from two adjacent stations at three depths in mangrove swamps on Techeng Island in Zhanjiang, China **(Supplementary Fig. 1 and Supplementary Table 1)**. Metagenomic *de novo* assembly and binning generated the reconstruction of over 112 archaeal genomes (> 50% complete). Among them, 22 genomes belonged to the Asgard superphylum. Partial high-quality Asgard genomes are shown in **Supplementary Table 2**. Bayesian and maximum likelihood phylogenic trees were reconstructed using 56 concatenated ribosomal proteins. The phylogenetic analyses revealed eight bins representing a novel lineage of the Asgard archaea **(Fig. 1a)**. These bins are located in distinct lineages and form a distantly related cluster with the Odinarchaeota in a phylogenetic tree with high bootstrap support **(Fig. 1a)**. They had a bin size ranging from 1.86 to 5.10 Mbp and a 43.1–48.9% mean GC content **(Table 1)**. Genome completeness of the bins ranged from 73.3% to 92.7% and there was almost no contamination of other genome fragments detected. These bins were recovered from top-layer (0.15–0.2 m) and mid-layer (0.4–0.45 m) samples of mangrove swamp sediment. For the three bins with > 85% completeness, their relative abundance in sequencing data from the six sediment samples ranged from 0.015% to 0.48% compared with the total raw reads **(Supplementary Table 10)**.

**Fig. 1.**
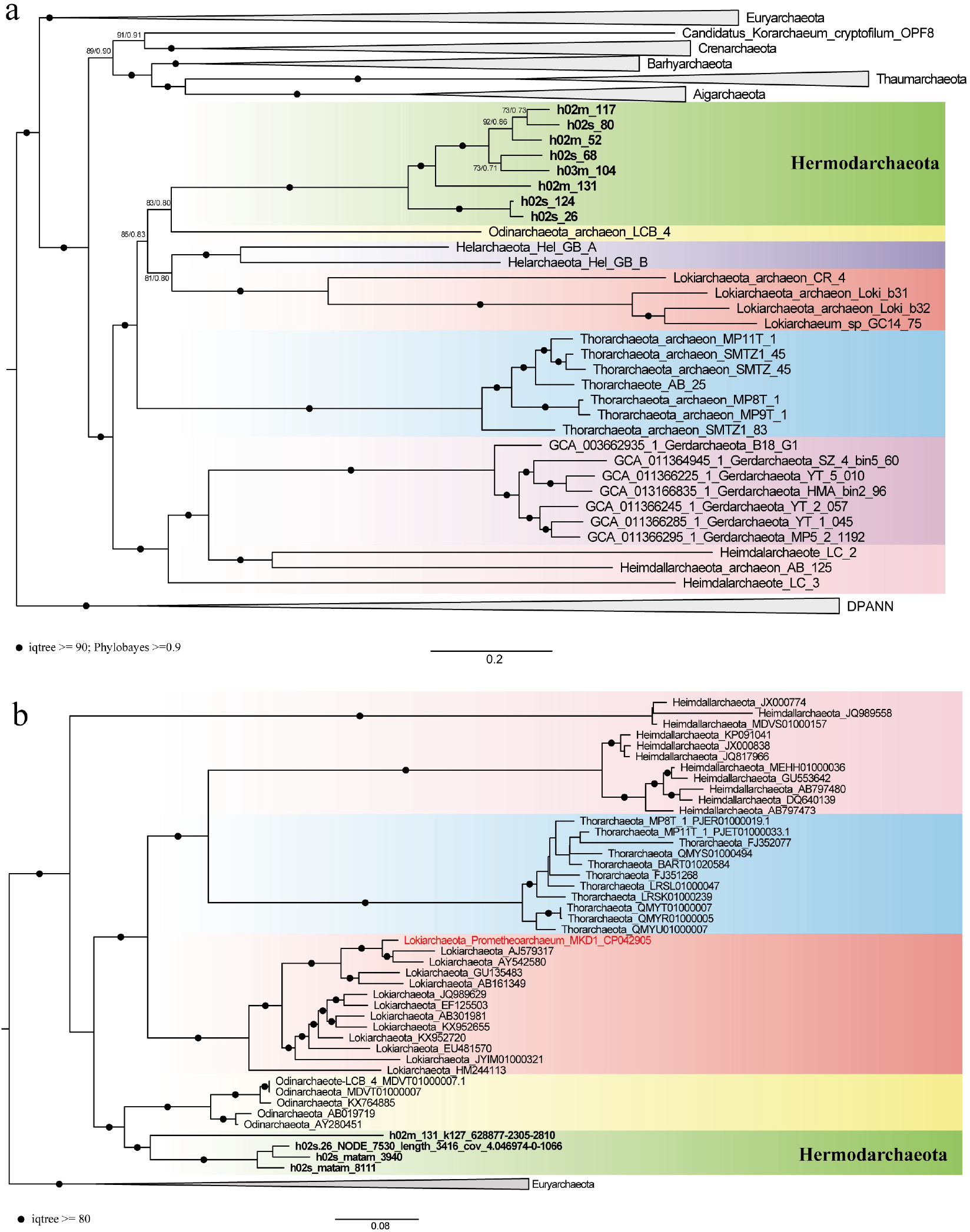
Phylogenetic placement of Hermodarchaeota within the Asgard archaea superphylum. **a** Phylogenomic tree of 56 concatenated ribosomal proteins reconstructed using IQtree with the LG+C60+F+G+PMSF model and PhyloBayes with the LG+CAT model. Nodes with ultrafast bootstrap values ≥ 90 and posterior probability ≥ 0.90 are indicated by black circles. **b** Maximum-likelihood phylogenetic tree of 16S rRNA gene sequences belonging to Hermodarchaeota were inferred using IQtree with GTR+F+I+G4 model. Black circles indicate bootstrap values ≥ 80.

**Table 1.** Genomic features of Hermodarchaeota bins

| Bin ID | h02s_80 | h02m_131 | h02s_68 | h02s_124 | h02m_52 | h02m_117 | h03m_104 | h02s_26 |
| --- | --- | --- | --- | --- | --- | --- | --- | --- |
| Completeness (%) | 92.67 | 89.5 | 86.22 | 73.33 | 77.46 | 74.69 | 78.04 | 76.42 |
| Contamination (%) | 0 | 1.87 | 0.47 | 0 | 0 | 0 | 0 | 1.94 |
| Strain heterogeneity (%) | 0 | 0 | 0 | 0 | 0 | 0 | 0 | 0 |
| Number of coding genes | 3636 | 3895 | 4833 | 2418 | 2582 | 1785 | 2561 | 2508 |
| GC content (%) | 44.54 | 43.21 | 43.91 | 48.97 | 44.52 | 44.71 | 43.05 | 48.74 |
| Contig/scaffold number | 267 | 863 | 903 | 338 | 357 | 269 | 469 | 677 |
| Estimated genome size (Mbp) | 3.76 | 4.22 | 5.10 | 2.45 | 2.68 | 1.86 | 2.66 | 2.53 |
| N50 (bp) | 20636 | 6335 | 6054 | 8604 | 11896 | 8731 | 6995 | 4119 |
| Longest contig/scaffold (bp) | 99342 | 54136 | 46141 | 48037 | 68536 | 36381 | 32924 | 28150 |

Genome completeness, contamination, and heterogeneity were estimated using CheckM<sup>43</sup>.

Two 16S rRNA genes were identified from bins of h02s_26 and h02m_131, respectively **(Supplementary Table 3)**. By blasting against all 16S rRNA gene sequences from the six samples, an additional two 16S rRNA genes were found to have more than 95% identity with that of h02s_26 **(Supplementary Tables 3 and Supplementary Table 11)**. Phylogenetic analyses revealed that the four 16S rRNA gene sequences formed a phylogenetically distinct group from the Odinarchaeota, and their position was similar to that of the above-mentioned eight draft genomic bins in the phylogenetic tree inferred by concatenated ribosomal proteins **(Fig. 1b)**. The 16S rRNA gene sequences of h02s_26 and h02m_131 showed a phylum level divergence with a DNA identity of 72.9–83.7% when compared with other Asgard archaeal 16S rRNA gene sequences (83.7% and 79.9% sequence similarity to that of Odinarchaeota, respectively) (**Supplementary Table 11**). Here, we propose *Candidatus* “Hermodarchaeota” as the name of this new group, after Hermod, the son of the god Odin in Norse mythology. An analysis of the average amino acid identity (AAI) revealed that these genomes had relatively low amino acid identity compared with other Asgard archaea (41.12–47.48%) **(Supplementary Fig. 3)**, further supporting their clustering as a separate phylum [58]. Although Hermodarchaeota bins formed a sister group with Odinarchaeota, they were markedly different from Odinarchaeota-LCB_4 in terms of total gene number and genome size. In addition, Hermodarchaeota bins possessed a suit of eukaryotic signature proteins (ESPs) that have been identified in other Asgard archaea **(Supplementary Table 12)**.

### Metabolic reconstruction of Hermodarchaeota

This study focused on the proteomes of three Hermodarchaeota bins with high completeness: h02s_80, h02m_131, and h02s_68. Similar to Lokiarchaeota and Thorarchaeota [55], metabolic analysis of Hermodarchaeota uncovered the presence of genes involved in the complete WLP (**Fig. 2**, **Supplementary Table 13**). The WLP is traditionally connected with methanogenesis in Archaea. However, all Hermodarchaeota genomes lacked genes that encode MCR and genes of key subunits that encode Na^+^-translocating methyl-THMPT:coenzyme M methyltransferase (MTR). Therefore, Hermodarchaeota are unable to perform hydrogenotrophic CO_2_-reducing methanogenesis. Each Hermodarchaeota genome contained three to five copies of gene encoding trimethylamine methyltransferases (mtt) and corresponding corrinoid proteins (mttc), as well as one to two copies of gene for methylcobamide: CoM methyltransferase (mtbA) and two to four copies of *mtrH* subunits. These genes may be sufficient for the production of methyltetrahydromethanopterin (methyl-H4MPT) from trimethylamine directly [59] or via methyl-coenzyme M [60] for further synthesis of Acetyl-coenzyme A (CoA). Metatranscriptome analyses showed that the *mtt*, *mttc*, and *mtbA* genes were expressed in mangrove sediments, supporting the transformation from trimethylamine to methyl-coenzyme M (**Fig. 2**, **Supplementary Tables 6-8)**.

**Fig. 2.**
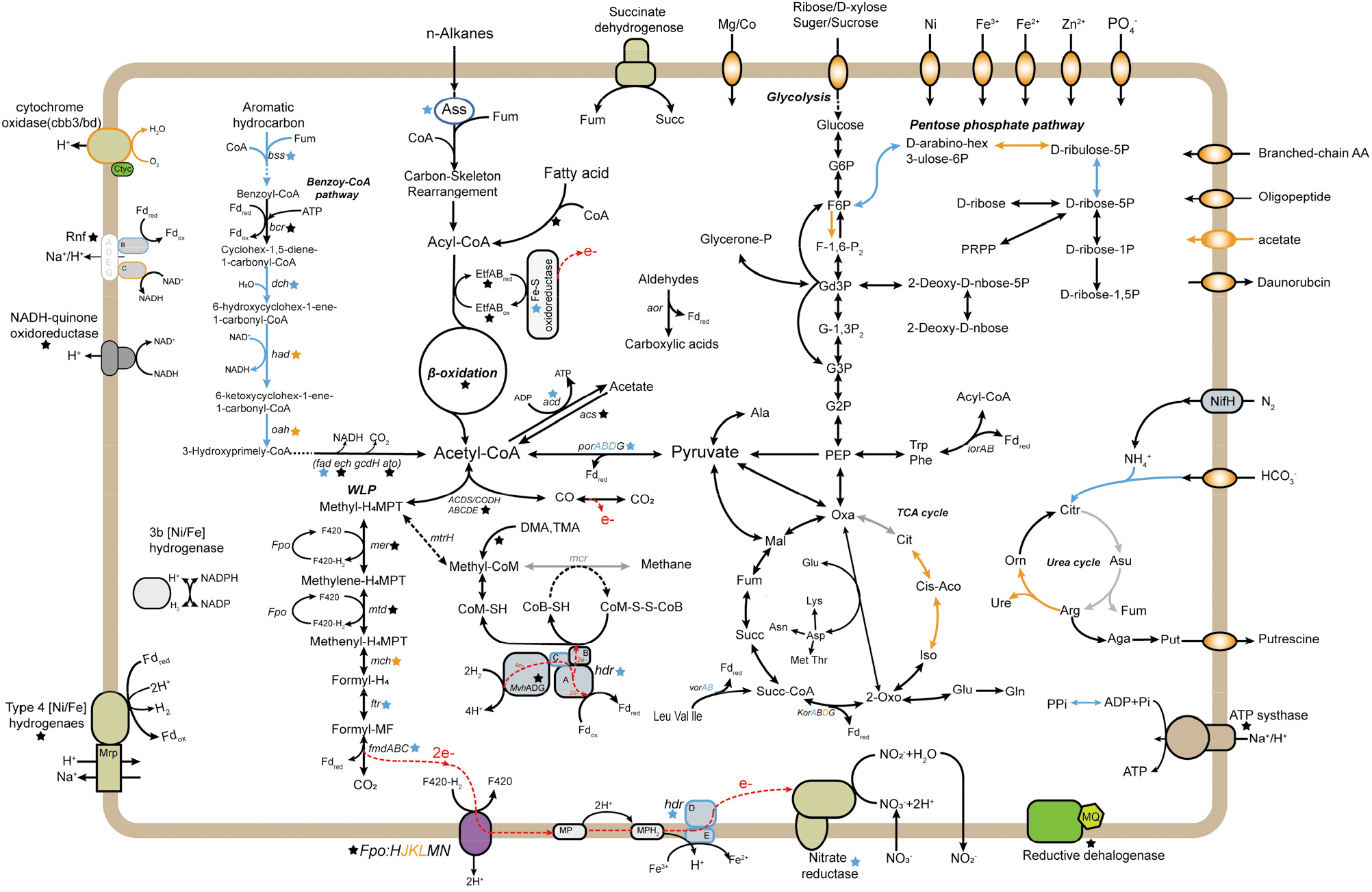
Key metabolic pathways in the three genomes of Hermodarchaeota: h02s_80, h02m_131, and h02s_68. Genes and pathways found in three bins (black), only found in two bins (blue), only found in one bin (orange), or missing from three bins (gray) are indicated. Red dotted lines represent electron flow. Genes related to the pathways presented in this figure are given in **Supplementary Table 13.** Gene or pathway names with an asterisk indicate that their transcripts have been detected in metatranscriptomes from mangrove wetlands **(Supplementary Tables 6-8)**. WLP, Wood-Ljungdahl pathway; TCA, tricarboxylic acid. DMA, dimethylamine; TMA, trimethylamine; MP, methanophenazine; Fd, ferredoxin.

In h02m_131 and h02s_68 genomes, three homologs of alkylsuccinate synthase (Ass) or benzylsuccinate synthase (Bss) were identified (**Fig. 2**, **Supplementary Table 13**). Their amino acid sequences exhibited 29–33% identity with AssA1 (ABH11460) from *D. alkenivorans* strain AK-01 (bit score: 262–312; e-values: e-98–7e-79) and 30–32% identity with BssA (YP158060) from *A. aromaticum* EbN1 (bit score: 234–294; e-values: 3e-90 – e-68), but only had 24–27% identity with the pyruvate formate lyase Pfl (NP415423) from *Escherichia coli* (bit score: 114–165; e-values: 4e-46–3e-29) **(Supplementary Table 14)**. Furthermore, higher similarity was observed between Ass/Bss of Hermodarchaeota and PflD (AAB89800) of *Archaeoglobus fulgidus* (identity: 33–38%; bit score: 336–463). The PflD has been suggested to possess an Ass activity in *A. fulgidus* [6]. The multiple sequence alignment revealed that, similar to other AssA and BssA, the three homologs in Hermodarchaeota harbor only one conserved cysteine which is used to receive the radical from the glycyl residue and initiate the reaction, whereas pyruvate formate lyases, such as Pfl from *E. coli*, possess two neighbouring conserved cysteines in the region [6] **(Fig. 3; Supplementary Fig. 4)**. Based on the 11 known archaeal sequences **(Fig. 3)**, AssA/BssA in archaea tends to substitute the first cysteine with glycine at the conserved region for PFLs. A phylogenetic analysis was performed using Hermodarchaeota Ass or Bss sequences and their closest homologs **(Supplementary Fig. 5)**. This revealed that Ass/Bss encoded by Hermodarchaeota are not monophyletic, but intermixed with bacterial sequences, indicating that these Ass/Bss sequences were likely obtained through horizontal gene transfers from bacteria, similar to the pflD of *A. fulgidus* [6].

**Fig. 3.**
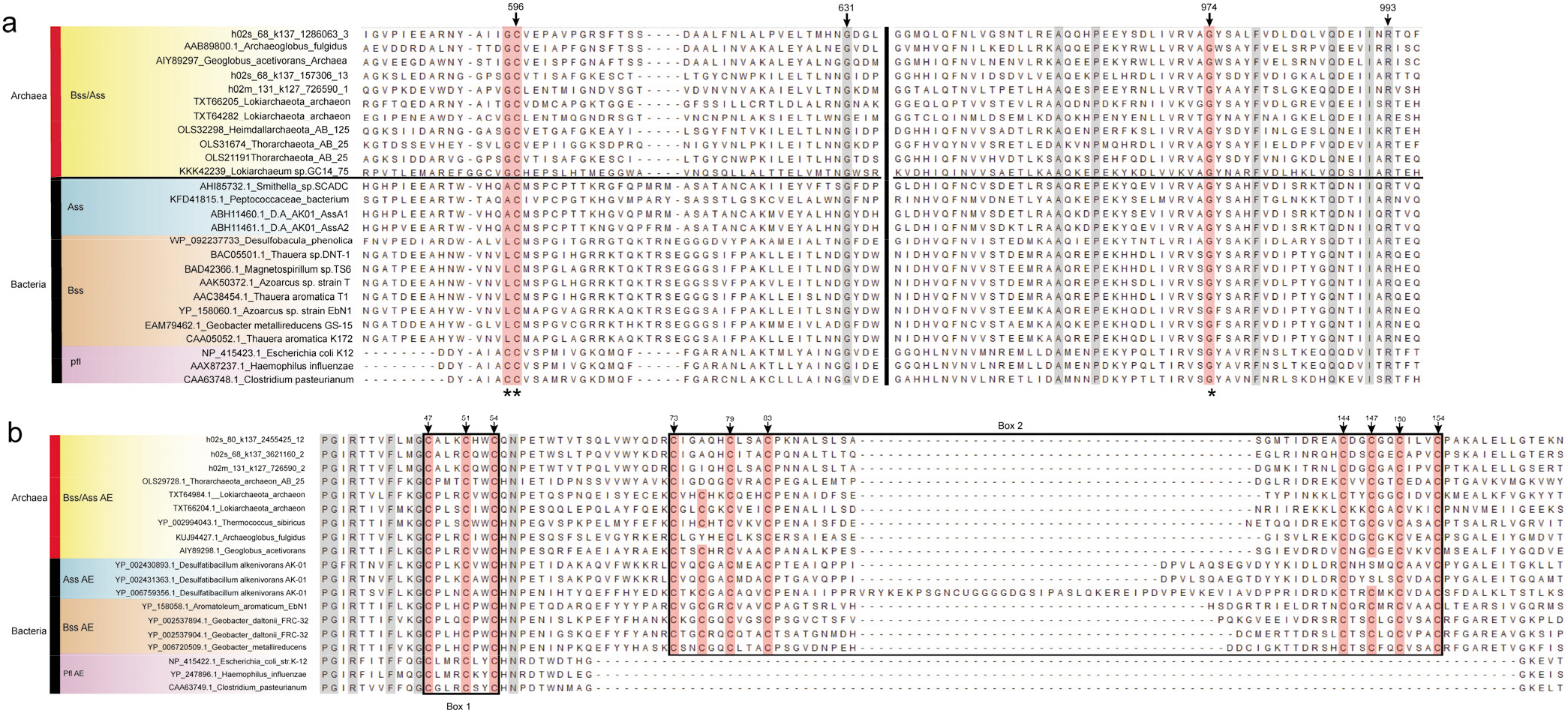
Partial sequence alignment of Hermodarchaeota alkylsuccinate synthase (Ass)/benzylsuccinate synthase (Bss) with known Ass/Bss and pyruvate formate lyase (pfl), and Hermodarchaeota Ass or Bss-activating enzyme (Ass/Bss AE) with known Ass/Bss and pyruvate formate lyase (pfl) AE. **a** Sequence comparison of Ass/Bss. The regions containing the conserved cysteine residue (**, shaded in red) and the conserved glycine residue (*, shaded in red) are presented. Vertical black line in the middle of sequences represents truncated section. Complete sequence alignments are given in **Supplementary Fig. 4**. Conserved residues are shaded in color. D. A_ AK-01, *D. alkenivorans* strain AK-01. **b** Sequence comparison of Ass/Bss AE. Boxes 1 and 2 correspond to the CxxxCxxC sequence motif and two cysteine-rich regions, respectively, and they are involved FeS cluster binding. Conserved residues are shaded in color. Complete sequence alignments are provided in **Supplementary Fig. 6.**

In addition to *Ass/Bss* genes, each Hermodarchaeota genome contained one gene encoding the Ass or Bss-activating enzyme (Ass/Bss AE), which is needed for Ass/Bss. A Blastp search found that they had greater similarity with AssD2 (YP_002431363) and AssD2’ (YP_002429341) from *D. alkenivorans* strain AK-01, PflC (KUJ94427) of *A. fulgidus*, and BssD (CAA05050) from *T. aromatica* K172, compared to the PflA-activating enzyme (NP_415422) of *E. coli* **(Supplementary Table 15)**. The N-terminal regions of Hermodarchaeota Ass/Bss AEs contained a CxxxCxxC sequence motif (box 1) and two cysteine-rich regions (box 2) **(Fig. 3, Supplementary Fig. 6).** Box1 is necessary for the Fe-S cluster of SAM-radical enzymes [61] while box 2 is involved in Fe-S cluster binding, which is unique to Ass/Bss AE and not found in pyruvate formate lyase-activating enzymes [6]. Generally, these data indicate that Hermodarchaeota possess Ass or Bss and activating enzymes. Furthermore, their corresponding transcripts were frequently detected in metatranscriptomes from mangrove sediments (**Fig. 2**, **Supplementary Tables 6-8)**, suggesting that these microorganisms are transcriptionally active in the anaerobic degradation of n-alkanes or aromatics via the addition to fumarate, producing alkyl-or benzyl-substituted succinates.

Once alkanes are activated, the alkyl-substituted succinates formed will be subjected to thioesterification, carbon-skeleton rearrangement, and decarboxylation [62]. At present, the genes involved in these reactions remain unclear. In *D. alkenivorans* strain AK-01, it is postulated that these steps were catalyzed by acyl-CoA synthetase (ligase) (AMP-forming), methylmalonyl-CoA mutase, and methylmalony-CoA carboxyltransferase [62]. The genes for all of these were present in each Hermodarchaeota genome **(Supplementary Table 13)** and were highly expressed in metatranscriptomes (**Supplementary Tables 6-8)**. Subsequently, acyl-CoA produced from alkane oxidation can be oxidized to acetyl-CoA by related enzymes of the beta-oxidation including acyl-CoA dehydrogenase (Acd), enoyl-CoA hydratase (Ech), 3-hydroxyacyl-CoA dehydrogenase (Hadh), and acetyl-CoA acyltransferase (Fad), and the genes encoding these enzymes have been identified in Hermodarchaeota genomes **(Supplementary Table 13)** and their corresponding transcripts were present and abundant in metatranscriptomes (**Supplementary Tables 6-8)**. In addition, each Hermodarchaeota genome contained genes for 10-19 Acd, two to three electron transfer flavoprotein complexes (ETF), and one to four FeS oxidoreductases **(Supplementary Table 13)**; transcripts for ETF and FeS oxidoreductase were identified (**Supplementary Tables 6-8)**. This could produce reduced ferredoxin or NADH by electron bifurcation in the ACD/ETF complex for anabolism (**Fig. 2**) [63, 64]. The genes for acetyl-CoA decarbonylase/synthase:CO dehydrogenase complex (ACDS/CODH) were identified in Hermodarchaeota **(Supplementary Table 13)** and their transcripts were frequently detected in metatranscriptomes (**Supplementary Tables 6-8)**; these are key enzymes in the metabolism of acetyl-CoA from beta-oxidation. This suggests that acetyl-CoA can be further oxidized into CO_2_ and yield reduced ferredoxin via the oxidative WLP as previously shown for butane oxidation in *Candidatus Syntrophoarchaeum* [65] **(Fig. 2)**.

For anaerobic oxidation of aromatic hydrocarbons, the first intermediate formed by the addition of fumarate, benzylsuccinate, is further oxidized to benzoyl-CoA, which is regarded as a primary aromatic intermediate in the anaerobic oxidation of plentiful aromatic hydrocarbons [66]. Hermodarchaeota genomes contained almost all genes found in the benzoyl-CoA pathway **(Fig. 2; Supplementary Table 13)**. Next in this pathway, the conversion from benzoyl-CoA to 3-hydroxypimelyl-CoA, is catalyzed by four key enzymes in *T. aromatica* including benzoyl-CoA reductase (Bcr), cyclohexa-1,5-dienecarbonyl-CoA hydratase (Dch), 6-hydroxycylohex-1-ene-1-carboxyl-CoA dehydrogenase (Had), and 6-oxocyclohex-1-ene-1-carbonyl-CoA hydrolase (Oah) [67], of which all genes were identified in h02s_80 and h02s_68 genomes (**Supplementary Table 13)**, and their corresponding transcripts were detected in metatranscriptomes (**Supplementary Tables 6-8)**. In addition, the genes encoding four subunits of Bcr in h02s_80 were found colocated in a contig with genes for an BzdV protein, a ferredoxin, a Dch, a Had, a NADP-dependent oxidoreductase, and two anaerobic benzoate catabolism transcriptional regulators **(Supplementary Fig. 7).** The arrangement of the gene cluster for benzoate catabolism was analogous to that in *T. aromatica* [42]. This indicates that ferredoxin probably acts as electron donor for the reduction of benzoyl-CoA in Hermodarchaeota while NADP-dependent oxidoreductase regenerates reduced ferredoxin as previously shown for *Azoarcus evansii* [68]. Subsequently, 3-hydroxypimelyl-CoA will be further oxidized to acetyl-CoA by the related enzymes via beta-oxidation including Hadh, Fad, glutaryl-CoA dehydrogenase (GcdH), glutaconyl-CoA decarboxylase (GcdA), Ech, 3-hydroxybutyryl-CoA dehydrogenase (PaaH), and acetyl-CoA acetyltransferase (AtoB). The genes for all these enzymes have been detected in Hermodarchaeota **(Fig. 2; Supplementary Table 13)**. Collectively, the data suggest that Hermodarchaeota are capable of using various aromatic hydrocarbons or alkanes as carbon and energy sources.

Compared to other Asgard archaea [55], Hermodarchaeota genomes possess more sophisticated energy-conserving complexes, including seven subunits of the F420H2 dehydrogenase (encoded by *fpo*), 11 subunits of NADH-quinone oxidoreductase, flavoprotein and FeS subunit of succinate dehydrogenase/fumarate reductase, group 4 [NiFe]-hydrogenase, the B and C subunits of the Rnf complex, and V/A-type adenosine triphosphate (ATP) synthase **(Fig. 2; Supplementary Fig. 8; Supplementary Table 13)**. Transcripts for partial subunits of these complexes were identified in metatranscriptomes (**Supplementary Tables 6-8)**. This suggests that these microorganisms can couple electron transfer with proton translocation across cytoplamic membrane and further generate an electrochemical proton gradient that drives ATP synthesis. Of note, two copies of genes that encode D and E subunits of CoB-CoM heterodisulfide reductase (Hdr) were identified in h02s_80 and h02s_68 genomes **(Fig. 2)**, which are absent in most methanogens and other Asgard archaea, but found solely in the *Methanosarcinales* [55, 69]. HdrED is an integral membrane complex, it accepts electrons from reduced methanophenazine (MPH2), and assists in the production of a chemiosmotic gradient across the cell membrane **(Fig. 2)**. The sophisticated energy-conserving system in Hermodarchaeota may enable better adaptability in the exploitation of various substrates for energy. In addition, *hdrABC* and *mvhADG* for methyl-viologen-reducing hydrogenase were also detected **(Fig. 2)**, suggesting that the complex possibly leads to co-reduction of CoM-S-S-CoB heterodisulfide and ferredoxin by electron bifurcation mechanism as shown previously for *Methanosarcina acetivorans* [69]. Owing to an absence of MCR, the reaction forming a heterodisulfide of CoM and CoB by MCR is not present in Hermodarchaeota. Although Hermodarchaeota contained genes for cytoplasmic fumarate reductase **(Supplementary Table 13)**, which has been shown to reduce fumarate using CoM-S-H and CoB-S-H in *Methanobacterium thermoautotrophicum* [70], the reaction is not accompanied by energy conservation. Thereby, like most Asgard archaea, it remains to be determined whether the cycle of CoM-S-S-CoB is operative in Hermodarchaeota.

In addition to metabolic pathways for aromatic hydrocarbons and alkanes, Hermodarchaeota genomes also contain multiple peptidases, aminopeptidases, carboxypeptidases, and amino acid and oligopeptide transporters (**Supplementary Fig. 9)**, suggesting that these microorganisms can utilize peptides and proteins as their sources of carbon and nitrogen. Similar to the peptide fermentation of *Pyrococcus furiosus* [71], amino acids hydrolyzed by these peptidases can be oxidatively deaminated by glutamate dehydrogenase (*gdh*), aspartate aminotransferases (*aspC*), 2-oxoglutarate ferredoxin oxidoreductase *(kor*), 2-ketoisovalerate ferredoxin oxidoreductase (*vor*), indolepyruvate ferredoxin oxidoreductase (*ior*), and pyruvate ferredoxin oxidoreductase (*por*) to generate acetyl-CoA and reduced ferredoxin **(Fig. 2; Supplementary Table 13)**. Furthermore, the aliphatic and aromatic aldehydes produced by these oxidoreductases can be oxidized to carboxylic acids by multiple aldehyde ferredoxin oxidoreductases, producing reduced ferredoxin **(Fig. 2)** [72]. The resulting acetyl-CoA is further catalyzed by acetyl-CoA synthetase to produce acetate (ADP-forming) **(Fig. 2)**, with concomitant formation of ATP [73]. In addition, similar to other Asgard archaea [8, 55], Hermodarchaeota may be capable of using carbohydrates as carbon or energy sources because they possess sugar transporters, sucrose transporters, and various carbohydrate-active enzymes **(Fig. 2, Supplementary Fig. 9)**. The resulting glucose can then be metabolized to generate intermediates for anabolism via glycolysis and an incomplete citric acid cycle, along with the formation of ATP and reduced nicotinamide adenine dinucleotide (NADH). Subsequently, the NADH is assumed to be oxidized by NADH-quinone oxidoreductase (complex I) to create a transmembrane proton gradient, or be applied to reduce ferredoxin via the Rnf complex (**Fig. 2**).

Proper electron acceptors are pivotal to successful degradation of aromatic compounds and alkanes. We did not identify any dissimilatory sulfite reductase (Dsr) or anaerobic sulfite reductase (Asr) in Hermodarchaeota genomes, indicating that they cannot perform dissimilatory sulfate reduction. However, Hermodarchaeota genomes contain genes encoding nitrate reductase, nitrate transporter NrtD and NrtB **(Fig. 2; Supplementary Table 13)**. Phylogenetic analysis showed that nitrate reductases of Hermodarchaeota formed two distinct clusters and were more closely related to the Nap/Nas clade from bacteria compared with the Nar clade **(Supplementary Fig. 10)**. Furthermore, motif analysis revealed that nitrate reductases of Hermodarchaeota might be homologues of periplasmic nitrate reductase (Nap) **(Supplementary Fig. 11)**. Transcripts for nitrate reductase were detected in metatranscriptomes (**Supplementary Tables 6-8)**. The results suggest that members of Hermodarchaeota are capable of utilizing nitrate as an electron acceptor during the oxidation of aromatic hydrocarbons as shown previously in denitrifying bacterium *Thauera aromatica* [74]. In addition to nitrate, Fe (III) has been found to serve as the sole electron acceptor of hyperthermophilic archaeon *Ferroglobus placidus* when oxidizing benzoate and phenol. In *Methanosarcina acetivorans*, electrons can be channeled to Fe (III) via HdrDE complex, which drives methane oxidation [75]. Thus, it is likely that Hermodarchaeota can couple oxidation of aromatic hydrocarbons with reduction of Fe (III) through the HdrDE complex **(Fig. 2)**. Similar to Lokiarchaeota and Thorarchaeota, identification of reductive dehalogenase in Hermodarchaeota genomes implicates that these microorganisms are also capable of performing organohalide respiration using chlorinated ethenes/ethanes **(Supplementary Fig. 12)**. In addition, a cytochrome bd-type quinol oxidase and a *cbb3-type* cytochrome c oxidase, which support aerobic respiration [76], were detected in h02m_131 and h02s_68 genomes (**Supplementary Table 13)**, respectively, indicating that some members of Hermodarchaeota possibly utilize oxygen as terminal electron acceptors under oxic environments, similar to Heimdallarchaeota of the Asgard archaea [55]. We could not identify any pilus and extracellular cytochromes that mediate electron transfer across species, as shown previously in *Ca. S. butanivorans* and ANME archaea [65, 77]. Therefore, it is unclear whether a syntrophic partner organism receiving reducing equivalents exists for members of Hermodarchaeota.

### Environmental distribution

To investigate the distribution and diversity of Ass/Bss sequences from archaea, *Ass/Bss* gene sequences from Hermodarchaeota were used to identify homologs in the six mangrove sediment samples in this study and across 1,000 publicly available metagenomes from around the world. A total of 394 nearly full-length *Ass/Bss* genes with archaeal GC motif were recovered from the six mangrove sediment samples (Techeng Island, China) (**Supplementary Table 16).** These sequences were divided into 14 clusters and exhibited an extraordinarily high genetic diversity **(Fig. 4a)**, likely suggesting that they may be derived from phylogenetically diverse archaea. Hermodarchaeota *Ass/Bss*-like gene fragments were also detected in various marine and freshwater environments, including marine bay sediments (Fagans, Australia), Guaymas Basin sediments enriched in hydrocarbon seeps (Gulf of California, USA), hot spring sediments (California, USA), deep-sea sediments with petroleum seeps (Eastern Gulf of Mexico), mangrove sediments (Yunxiao, China), Towuti Lake sediments (South Sulawesi, Indonesia), and formation water in coal beds (Qinshui Basin, China) **(Fig. 4b, Supplementary Table 9)**. Furthermore, in these environments, a considerable number of 16S rRNA gene fragments were found to have high sequence similarity (≥ 90% or 95% identity) to that of Hermodarchaeota **(Fig. 4b, Supplementary Table 9)**. The homologs of Ass/Bss were also identified in the thermophilic pure archaeon *A. fulgidus* isolated from a submarine hot vent [6] as well as in composite genomes of Thorarchaeota and Lokiarchaeota from deep seabed petroleum seeps [78]. These results suggest that members of Hermodarchaeota, and other archaea capable of performing oxidation of alkanes and aromatic hydrocarbons through addition to fumarate, may be ubiquitous in nature.

**Fig. 4.**
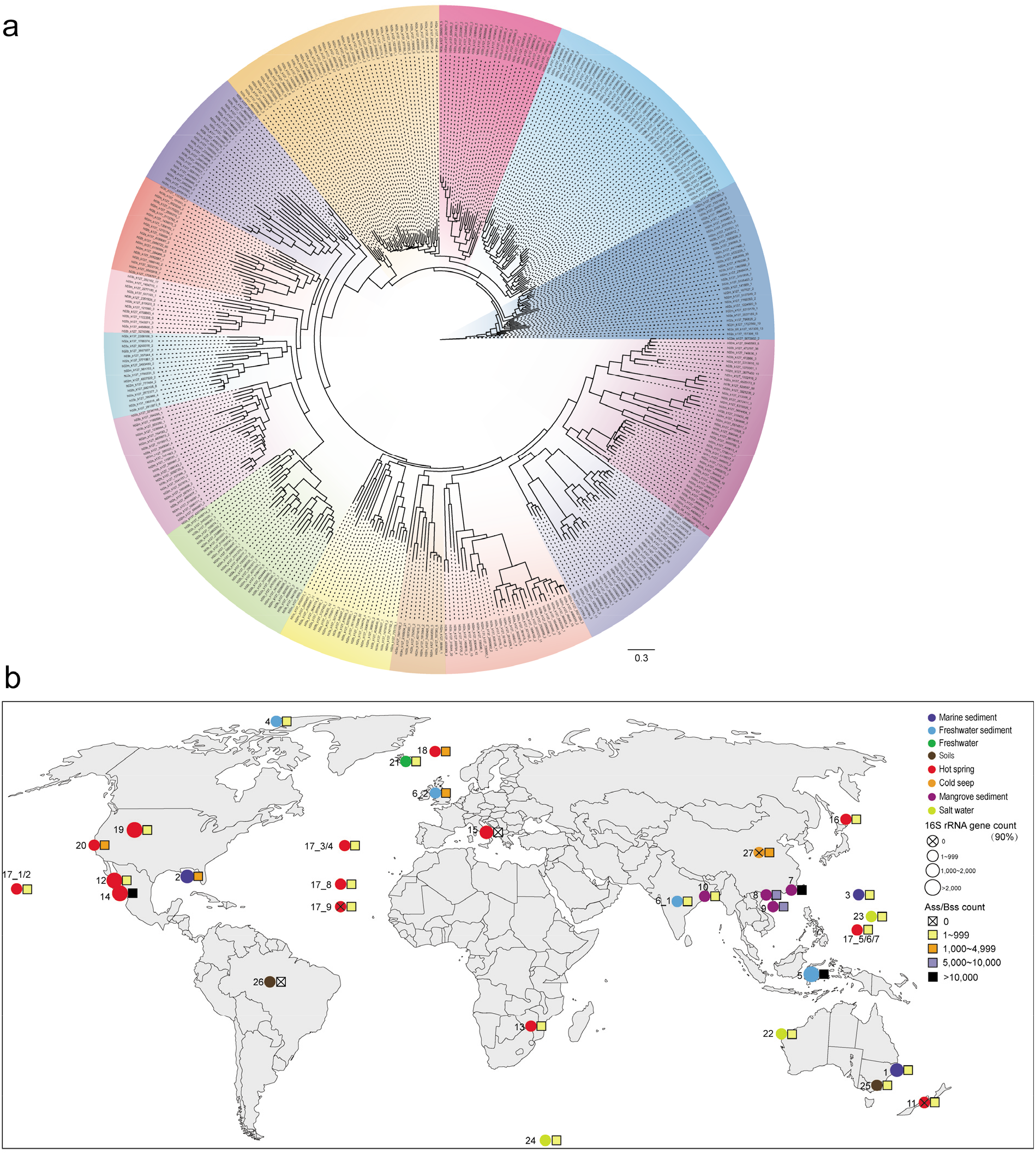
Maximum-likelihood tree of homologs of Hermodarchaeota alkyl/benzyl-succinate synthase (Ass/Bss) identified in metagenomes from the six mangrove sediments on Techeng Island, reconstructed using IQtree with the LG+F+R10 model (a), as well as global distribution of homologs of Hermodarchaeota Ass/Bss and 16S rRNA genes identified in metagenomes from various environments (b). **a** Features of these Ass/Bss sequences for Blast results are given in **Supplementary Table 16**; Sequence alignments show that they all contain “GC” motif for archaeal Ass/Bss. **b** Circles with different colors represent metagenomes from different habitats; the size of the circles indicate the range of 16S rRNA gene count; the range of Ass/Bss gene count are indicated using squares with different colors; the numbers in the figure correspond to the ID in **Supplementary Table 9**; detailed information is provided in **Supplementary Table 9**.

In addition to utilization of alkane, Hermodarchaeota is able to perform anaerobic oxidation of aromatic hydrocarbon via addition to fumarate coupling with the benzoyl-CoA degradation pathway. The three subunits of the key enzyme Bcr for the benzoyl-CoA pathway are highly related to those of ATP-consuming class I Bcr of Anaerolineales bacterium of Chloroflexi (Supplementary Fig. 13), suggesting occurrence of horizontal gene transfers between Hermodarchaeota and Chloroflexi. The *bcr* genes and genes for downstream transformation were also identified in composite genomes of Thermoplasmata and Bathyarchaeota from deep-sea sediments with petroleum seeps [78]. In addition, a new pathway for anaerobic oxidation of short-chain hydrocarbon via alkyl-coenzyme M formation has been proposed in Bathyarchaeota, Hadesarchaeota, *Ca*. Syntrophoarchaeum, and Helarchaeota [7, 8]. These results demonstrate that metabolic processes of hydrocarbons in archaea may be more complicated than thought before. Such complexity likely suggests that utilization of hydrocarbons by archaea may have existed for a long time in the earth. The discovery of Hermodarchaeota and its ubiquitous distribution expands the domain of archaea and has crucial significance for understanding of the ecological functions and evolutionary history of the mysterious Asgard archaea.

## Acknowledgements

This study was supported by the National Natural Science Foundation of China (41971125 and 41725002), the Guangdong Natural Science Foundation (2018A030313164), and the Innovation Study Project of East China Normal University. We thank Silvia E. Newell for editing this manuscript.

## Author contributions

H.P.D, L.J.H, M.L, J.W.Z and M.L conceived the study. J.W.Z and Y.F.O recovered genomes from metagenomes and analyzed these genomic data. J.W.Z, H.P.D, Y.L.Z and P.H performed metatranscriptome analyses. X.L. A.G.Y. Y.L and D.M.W performed analyses of phylogenies and environmental distribution of genes. H.P.D, L.J.H, M.L, and Y.L wrote the manuscript and supplementary information with contributions from all co-authors. All authors provided comments on the manuscript.

## Additional information

Supplementary information is available for this paper.

