## Supplemental Tables and Figures for "Novel Asgard archaea phylum Hermodarchaeota degrade alkanes and aromatics via alkyl/benzyl-succinate synthase and benzoyl-CoA pathway"

^1^State Key Laboratory of Estuarine and Coastal Research, East China Normal University, Shanghai 200062, China; ^2^School of Ocean and Meteorology, Guangdong Ocean University, Zhanjiang 524088, China; ^3^Shenzhen Key Laboratory of Marine Microbiome Engineering, Institute for Advanced Study, Shenzhen University, Shenzhen 518060, China; ^4^Key Laboratory of Geographic Information Science, Ministry of Education, East China Normal University, Shanghai 200061, China

**Table of contents**

**A. Supplementary Methods	2-3**

1. Sulfite reductase analyses

2. Sequence phylogeny of ribulose-1,5-bisphosphate carboxylase/oxygenase

3. Identification and phylogeny of Hermodarchaeota alkyl/benzyl-succinate synthase homologues

**B. Supplementary Discussion	3-8**

1. Carbon metabolism

2. System of energy conserving electron transport

3. Environmental distribution

**C. Supplementary Tables and Figures	9-36**

**Suppl. Tables 1-5, Tables 10-11, and 14-15	9-18**

**Legends for Suppl. Tables 6-9, 12-13 and 16**

**(provided in separate excel files)	19**

**Suppl. Figures 1-14	20-34**

**References	35-37**

**A. Supplementary Methods**

**1. Sulfite reductase analyses**

Identification of dissimilatory sulfite reductase (Dsr) and anaerobic sulfite reductase (Asr) genes, which are key enzymes for dissimilatory sulfate reduction, was performed by searching all predicted genes in Hermodarchaeota bins against *dsrA, dsrB, dsrD, asrA, asrB*, and *asrC* from TIGRfam [1] and Pfam [2] using hmmscan [3] (v.3.2.1). Motifs and conserved residues for DsrA, DsrB, DsrD, and AsrC proteins were analyzed by aligning the identified genes with known reference proteins [4]. Dozens of protein sequences were found to harbor Dsr or Asr domains. However, these sequences lacked strictly conserved binding sites for the siroheme-[4Fe4S] found in canonical DsrA, DsrB, and AsrC [4].

**2. Sequence phylogeny of ribulose-1,5-bisphosphate carboxylase/oxygenase**

Representative sequences of different forms of ribulose-1,5-bisphosphate carboxylase/oxygenase (Rubisco) reported previously by Tabita et al. [5], along with Rubisco sequences from Hermodarchaeota and other Asgard lineages, were aligned using MAFFT-L-INS-I [6] and trimmed using BMGE [7] (BLOSUM30 option). A maximum-likelihood phylogenic tree was inferred using IQtree [8] (v.1.6.12) under the LG+I+G4 model. Support values were calculated using 1,000 ultrafast bootstraps and SH-like approximate likelihood ratio test.

**3. Identification and phylogeny of Hermodarchaeota alkyl/benzyl-succinate synthase** **homologues**

Metagenomic sequences from the six mangrove sediments collected from Techeng Island were *de novo* assembled individually using MEGAHIT [9] with the following parameters: --k-min 27--k-max 127--k-step 10. Protein coding sequences in the resulting contigs were predicted using Prodigal [10] (v.2.6.3) using prodigal -a pep.fas -c cds.fa -i contigs.fa -p meta. Alkyl/benzyl-succinate synthase (Ass/Bss) sequences were identified by searching all predicted genes against Hermodarchaeota Ass/Bss sequences using Blastp [11] with an E-value < 1e-5. From the BLAST results, hits with a bit score ≥ 400 and length ≥ 650 amino acids were sorted and further filtered using CD-HIT [12] with a cut-off of 90% amino acid sequence identity. The motifs of Ass/Bss were analyzed following a previous method [13]. As a result, 394 sequences with archaeal “GC” motif were obtained from the six metagenomic samples. These sequences were aligned using MAFFT-L-INS-I [6]. The resulting alignments were trimmed using BMGE [7] (BLOSUM30 option). A maximum-likelihood phylogenic tree was reconstructed using IQtree [8] (v.1.6.12) under the LG+F+R10 model. Support values were calculated using 1,000 ultrafast bootstraps.

**B. Supplementary Discussion**

**1. Carbon metabolism**

Similar to other Asgard and Euryarchaeota archaea [14, 15], Hermodarchaeota possesses a modified Embden-Meyerhof pathway **(****Supplementary Table 13)**. It lacked canonical hexokinase catalyzing phosphorylation of glucose, but contained multiple sugar and carbohydrate kinases harboring the domain of the pfkB family carbohydrate kinase (PF00294), which have been found in other Asgard archaea. It is likely that these enzymes represent a new family of kinases capable of phosphorylating hexoses. We did not identify bifunctional ADP-dependent phosphofructokinase/glucokinases, which use ADP as the phosphoryl donor and have previously been identified in Heimdallarchaeota and several archaea of the Euryarchaeota [14, 15]. In addition, genes encoding a canonical glucose-6-phosphate isomerase were not detected in Hermodarchaeota; however, multiple genes for sugar phosphate isomerase/epimerase were present and may perform the same function **(Supplementary Table 13)**. Like several Asgard archaeal lineages, Hermodarchaeota contained genes encoding bifunctional archaeal fructose-1 6-bisphosphate aldolase (FBP)/phosphatase, with both FBP aldolase and FBP phosphatase activity [16], and canonical fructose-1, 6-bisphosphate phosphatase, suggesting alternative options in the gluconeogenesis reaction.

Members of Hermodarchaeota contained an incomplete tricarboxylic acid (TCA) cycle **(Fig. 2),** similar to Odinarchaeum and Lokiarchaeum [15]. The three bins of Hermodarchaeota all lacked genes for citrate synthase and citrate lyase while genes for aconitate hydratase or isocitrate dehydrogenase were only detected in one bin **(Fig. 2**, **Supplementary Table 13)**. This suggests that Hermodarchaeota is unable to oxidize acetyl-CoA derived from organic compounds into CO_2_ by the oxidative TCA cycle or to fix CO_2_ by the reductive TCA cycle. However, the incomplete TCA cycle may be able to mediate breakdown and synthesis of amino acids **(Fig. 2).**

Genes encoding all enzymes for the Wood–Ljungdahl pathway (WLP) were detected in each bin of Hermodarchaeota **(Fig. 2)**, similar to Thorarchaeota and Lokiarchaeum. The WLP consists of methyl and carbonyl branches [17]. In the carbonyl branch, a bifunctional and oxygen-sensitive enzyme complex, carbon monoxide dehydrogenase/acetyl-CoA synthase, consisting of five subunits, was identified in each genome of Hermodarchaeota **(Supplementary Table 13)**. The complex can catalyze both the reaction from CO, CoA, and methyl derived from the methyl branch to synthesize acetyl-CoA, and the reaction from acetyl-CoA to produce CO_2_ and methyl-H_4_MPT [18]. In methanogens, the methyl-H_4_MPT: coenzyme M methyltransferase (MTR) can transfer a methyl from H_4_MPT to coenzyme M (CoM) to form methyl-CoM coupling with translocation of Na^+^ across the membrane [19]. Subsequently, the methyl-CoM produced is converted to CH_4_ by methyl-coenzyme M reductase complex (MCR). The MTR complex consists of eight subunits [20]; however, only subunit H was found in Hermodarchaeota **(Fig. 2).** In addition, Hermodarchaeota lacked genes encoding all subunits of MCR. The feature is similar to all of the Asgard archaeal lineages except for Helarchaeota. This suggests that Hermodarchaeota is unable to perform methanogenesis. It is proposed that the presence of the archaeal WLP in the absence of MCR and MTR complexes might be the remnant of a previous combination with methanogenesis [19].

Several genes encoding 1,5-bisphosphate carboxylase/oxygenase (Rubisco) were identified in each genome of Hermodarchaeota. This enzyme is also present in all members of the Asgard archaea [15]. Four types of Rubisco are found in nature. Types I, II, and III catalyze the carboxylation and oxygenation of ribulose 1, 5-bisphosphate, which is the primary CO_2_ fixation reaction in the Calvin-Benson-Bassham cycle, whereas type IV does not catalyze either of these reactions [5]. Members of Hermodarchaeota contained nine genes for the large subunit of Rubisco. Phylogenetic analysis showed that six Rubisco proteins in Hermodarchaeota belonged to type IV and three belonged to archaeal type III (**Supplementary Fig. 14).** The archaeal type III Rubisco proteins have also been found in Helarchaeota, Lokiarchaeota, and Heimdallarchaeota [15] (**Supplementary Fig. 14)**. The real function of these type III Rubisco proteins in Asgard archaea requires further study.

**2. System of energy conserving electron transport**

Genes encoding 11 subunits of NADH-quinone oxidoreductase were identified in Hermodarchaeota including *nuoA, B, C, D, H, I, J, K, L, M,* and *N* (**Supplementary Table 13)**, each subunit had multiple copies in one bin*.* However, in other Asgard archaea, only Heimdallarchaeota and Thorarchaeota possess a multisubunit NADH-quinone oxidoreductase (3–5 subunits) [15]. This enzyme, also called complex I, is an energy conserving enzyme that transfers electrons from NADH to quinones, coupled with translocation of protons across the membrane. The complex I and group 4 hydrogenase showed a close relationship. A previous method was used to differentiate between them [21]. According to this method, the NADH-quinone oxidoreductase of Hermodarchaeota lacked subunits EFG while no CxxC motifs were found in its nuoD; therefore, it should belong to complex I-related enzymes, but not group 4 hydrogenases. The presence of complex I suggests that Hermodarchaeota has potential to use NADH derived from the Embden-Meyerhof-Parnas pathway to produce energy.

Seven subunits of F_420_H_2_ dehydrogenase (Fpo DHLKMNJ) were found in the genomes of Hermodarchaeota (**Supplementary Table 13).** In contrast, multi-subunit F_420_H_2_ dehydrogenase has not been found in other Asgard archaeal lineages [15]. In methanogenic archaeon, F_420_H_2_ dehydrogenase is a membrane-bound energy conserving enzyme that is coupled to proton translocation across the membrane by catalyzing F_420_H_2_-dependent reduction of methanophenazine [22]. It is assumed that F_420_H_2_ dehydrogenase may be responsible for oxidation of F_420_H_2_ produced in oxidative WLP **(Fig. 2).**

Similar to other Asgard archaea, several genes encoding the large subunit of group 3 and group 4 [NiFe]-hydrogenases were identified in each genome of Hermodarchaeota via HydDB and phylogenetic analyses (**Supplementary Fig. 8)**. The N-terminal and C-terminal region of these hydrogenase sequences harbor a conserved CxxC motif (**Supplementary Fig. 8),** which can bind the [NiFe] centers for H_2_ oxidation [23]. In phylogenetic trees, most hydrogenase sequences from Hermodarchaeota can form a small cluster with those of other Asgard archaea. However, five group 3 hydrogenase sequences from Hermodarchaeota formed a novel cluster within group 3b (**Supplementary Fig. 8)**. Hermodarchaeota contained both group 3b and 3c hydrogenases. In the hyperthermophilic archaeon, *Pyrococcus furiosus*, group 3b hydrogenase possesses a NADP/FAD binding domain and can use H_2_ to reduce NADP for other biosynthetic metabolism [24] **(Fig. 2)**. In methanogens, group 3c hydrogenase is a methyl-viologen-reducing hydrogenase (Mvh) comprising three subunits (ADG). The enzyme is associated with heterodisulfide reductase (hdrABC) in the reduction of CoM-S-S-CoB[24] **(Fig. 2)**. The two group 3c hydrogenases of Hermodarchaeota were annotated as methyl-viologen-reducing hydrogenase alpha subunit (arCOG01549) in the arCOG database (**Supplementary Table 13).** The results suggest that Hermodarchaeota may be able to use H_2_ as electron donor. Phylogenetic analysis showed that most of group 4 hydrogenases of Hermodarchaeota belonged to group 4g (**Supplementary Fig. 8)**, which is a H_2_-evolving membrane-bound hydrogenase. In *P. furiosus,* a 14-subunit membrane bound hydrogenase (MBH) oxidizes reduced ferredoxin generated by sugar fermentation and releases H_2_ coupled to translocation of sodium ion across the cell membrane [25]. It has been found that many subunits for MBH are homologous to subunits of complex I and Mrp H^+^/Na^+^ antiporter [26]. As mentioned above, many subunits in complex I for each genome of Hermodarchaeota had multiple copies in which some possessed the domain of the Mrp antiporter (such as nuoLMN) (IPR001750). It is likely that some subunit copies in complex I may be a part of the MBH complex in Hermodarchaeota. Thus, members of Hermodarchaeota may be capable of using MBH to establish a Na^+^ gradient for ATP synthesis **(Fig. 2).**

Each genome of Hermodarchaeota contained genes for reductive dehalogenase. These enzymes were closely related to reductive dehalogenase from Bathyarchaeota archaeon and Verstraetearchaeota archaeon (E-value: 2.88E-151 to 6.56E-151; bit score: 439), and contained a domain of reductive dehalogenase (IPR028894). Reductive dehalogenases perform dehalogenation in organohalide respiring bacteria that are responsible for the decomposition of many organohalide pollutants, such as polychlorinated biphenyls and dioxins [27, 28]. Phylogenetic analyses using a previous dataset with functionally characterized reductive dehalogenases [29] revealed that reductive dehalogenases of Hermodarchaeota were grouped with those of Lokiarchaeota, Thorarchaeota, and Heimdalarchaeota, as well as of *Dehalospirillum multivorans* PceA (O68252) and *Desulfitobacterium* sp. PCE-1 (**Supplementary Fig. 12),** suggesting that these Asgard archaea may be able to utilize chlorinated ethenes/ethanes as electron acceptors.

A total of 10 nitrate reductases were identified in the genomes of Hermodarchaeota using our method (see Methods section) (E-value: 3.39E-61 to 2E-154; bit score: 216 to 473) (**Supplementary Fig. 10)**. They all contained a molybdopterin oxidoreductase domain (IPR006656) and a CXXCXXXC motif (**Supplementary Fig. 10).** Prokaryotic nitrate reductases form two major clades, the membrane-associated prokaryotic nitrate reductase (Nar) clade, and the periplasmic nitrate reductase (Nap)/prokaryotic assimilatory nitrate reductase (Nas) clade [30]. It has been reported that all Nap sequences have an iron-sulfur cluster with a CXXCXXXC motif at the N-terminal region [30]. Phylogenetic analyses showed that nitrate reductases of Hermodarchaeota were grouped into a large cluster with the Nap/Nas clade but formed two distinct deep branches, suggesting that they may be more ancient than bacterial Nap/Nas. Thus, it is inferred that these nitrate reductases of Hermodarchaeota may be analogous in function to prokaryotic Nap. In some denitrifiers, Nap has been suggested to perform dissimilatory nitrate reduction [31]. It is likely that Hermodarchaeota may have potential to use nitrate as a terminal electron acceptor.

**3. Environmental distribution**

Although gene fragments homologous to Hermodarchaeota alkyl/benzyl-succinate synthase (Ass/Bss) were detected in a variety of environments, they were mainly found in high abundance in metagenomes from deep sea petroleum seepages, marine hydrothermal vents, hot springs, mangrove swamps, lake sediments, and formation water from coal seams, while only detected in low abundance in metagenomes from deep-sea water, Amazon forest soil, and freshwater wetland sediments (**Supplementary Table 9; Fig. 4).** This appears to be consistent with the content of a variety of alkanes or aromatic hydrocarbons contained in these environments. The presence of abundant Ass/Bss genes in many mangrove wetlands may be due to release of large amount of lignin-derived phenols and n-alkanes from mangrove tissues as well as aromatic compounds produced by lignin degradation [32]. The Hermodarchaeota 16S rRNA gene was identified in most of these environments, implicating ubiquity of members of Hermodarchaeota in nature. In certain habitats, the high abundance of Ass/Bss gene fragments corresponded to a high percentage of the Hermodarchaeota 16S rRNA gene (**Supplementary Table 9; Fig. 4)**, suggesting a substantial contribution of Hermodarchaeota to anaerobic degradation of alkanes or aromatic hydrocarbons. Furthermore, these Ass/Bss gene fragments may be derived from diverse archaeal lineages based on our study in mangrove wetlands **(Fig. 4a).** Thus, archaea capable of performing anaerobic degradation of alkanes or aromatic hydrocarbons through addition to fumarate may be more diverse in the environment than previously thought.

**Supplementary Table 1.** Sediment characteristics of sampling sites.

| **Samples** | **Location** | **Depth**  **(m)** | **pH** | **Eh**  **(mV)** | **DO**  **(mg L^-1^)** | **NH_4_^+^**  **(μg g^-1^)** | **NO_3_^-^**  **(μg g^-1^)** | **NO_2_^-^**  **(μg g^-1^)** |
| --- | --- | --- | --- | --- | --- | --- | --- | --- |
| h02s | H02, 110°26′45″, 21°10′27″ | 0.15-0.20 | 7.4 | -311.02 | ND | 10.39 | 0.59 | 0.14 |
| h02m |  | 0.40-0.45 | 7.1 | -230.73 | ND | 9.94 | 0.64 | 0.08 |
| h02b |  | 0.95-1.0 | 6.7 | -197.25 | ND | 9.85 | 0.58 | 0.08 |
| h03s | H03, 110°26′19.67″,  21°9′10.80″ | 0.15-0.20 | 7.4 | -254.35 | ND | 10.67 | 0.57 | 0.06 |
| h03m |  | 0.40-0.45 | 6.6 | -253.92 | ND | 10.36 | 0.56 | 0.04 |
| h03b |  | 0.95-1.0 | 6.3 | -167.44 | ND | 9.76 | 0.72 | 0.07 |

^*^Sediment ph, redox potential (Eh), and dissolved oxygen (DO) were in triplicate measured using a micromanipulator meter **system (Unisense,** Denmark) with a needle pH sensor, a needle Eh senor and a needle oxygen sensor, respectively. Sediment ammonium (NH_4_^+^), nitrate (NO_3_^–^) and nitrite (NO_2_^–^) were extracted by 2 mol L^-1^ KCl, and their concentrations were determined via flow injection analysis (Skalar Analytical SAN++, Netherlands). ND represents the values below the detection limit (0.05 μmol L^-1^).

**Supplementary Table 2.** Statistics of Asgard archaeal bins recovered from mangrove sediment samples

| **Bin ID** | **Assembly tool used** | **Completeness (%)** | **Contamination (%)** | **Strain heterogeneity (%)** | **NO. of gene** | **GC content (%)** | **NO. of Scaffold**  **/contigs** | **Genome Size**  **(Mbp)** | **Largest scaffold/contig**  **(bp)** | **Taxonomy** |
| --- | --- | --- | --- | --- | --- | --- | --- | --- | --- | --- |
| h02s_68 | Megahit | 86.22 | 0.47 | 0 | 4833 | 43.91 | 903 | 5.1 | 46141 | Hermodarchaeota |
| h02s_80 | Megahit | 92.67 | 0 | 0 | 3636 | 44.54 | 267 | 3.76 | 99342 | Hermodarchaeota |
| h02s_124 | Megahit | 73.33 | 0 | 0 | 2418 | 48.97 | 338 | 2.45 | 48037 | Hermodarchaeota |
| h02m_52 | Megahit | 77.46 | 0 | 0 | 2582 | 44.52 | 357 | 2.68 | 68536 | Hermodarchaeota |
| h02m_117 | Megahit | 74.69 | 0 | 0 | 1785 | 44.71 | 269 | 1.86 | 36381 | Hermodarchaeota |
| h02m_131 | Megahit | 89.5 | 1.87 | 0 | 3895 | 43.21 | 863 | 4.22 | 54136 | Hermodarchaeota |
| h03m_104 | Megahit | 78.04 | 0 | 0 | 2561 | 43.05 | 469 | 2.66 | 32924 | Hermodarchaeota |
| h02s_26 | metaSPAdes | 76.42 | 1.94 | 0 | 2508 | 48.74 | 677 | 2.53 | 28150 | Hermodarchaeota |
| h02s_84 | Megahit | 73.39 | 2.96 | 0 | 1722 | 38.15 | 442 | 1.78 | 11729 | Heimdallarchaeota |
| h03b_10 | Megahit | 79.16 | 2.49 | 0 | 1666 | 37.69 | 696 | 1.73 | 10281 | Heimdallarchaeota |
| h02m_144 | Megahit | 77.93 | 0.93 | 100 | 3393 | 29.81 | 640 | 3.55 | 56443 | Lokiarchaeota |
| h02m_142 | Megahit | 90.63 | 0 | 0 | 2749 | 42.7 | 306 | 2.66 | 69294 | Odinarchaeota |
| h02s_33 | Megahit | 60.85 | 0 | 0 | 2289 | 44.29 | 564 | 2.36 | 13267 | Thorarchaeota |

**Supplementary Table 3.** 16S rRNA genes of Hermodarchaeota.

| **ID** | **Samples** | **Assembly tool** | **Scaffold/Contig length** | **Position of 16S rRNA gene** | **Length of 16S rRNA gene** | **Bin of Hermodarchaeota** |
| --- | --- | --- | --- | --- | --- | --- |
| h02s.26_NODE_7530 | h02s | metaSPAdes | 3416 | 1-1066 | 1066 | h02s_26 |
| h02m_131_k127_628877 | h02m | Megahit | 2810 | 2305-2810 | 506 | h02m_131 |
| h02s_matam_3940 | h02s | MATAM | - | - | 1450 | - |
| h02s_matam_8111 | h02s | MATAM | - | - | 879 | - |

**Supplementary Table 4.** List of organisms used in phylogenetic analyses.

| **Domin** | **Taxonomic level 1** | **Taxonomic level 2** | **Genomes** | **Accession** | **Ribosomal tree** | **16S tree** |
| --- | --- | --- | --- | --- | --- | --- |
| Archaea | Asgard | Candidatus Heimdallarchaeota | Heimdallarchaeota archaeon AB_125 | MEHH01 | 1 | 1 |
| Archaea | Asgard | Candidatus Heimdallarchaeota | Heimdalarchaeote LC_2 | MDVR01 | 1 | 0 |
| Archaea | Asgard | Candidatus Heimdallarchaeota | Heimdalarchaeote LC_3 | MDVS01 | 1 | 1 |
| Archaea | Asgard | Candidatus Heimdallarchaeota | - | KP091041 | 0 | 1 |
| Archaea | Asgard | Candidatus Heimdallarchaeota | - | JX000838 | 0 | 1 |
| Archaea | Asgard | Candidatus Heimdallarchaeota | - | AB797480 | 0 | 1 |
| Archaea | Asgard | Candidatus Heimdallarchaeota | - | AB797473 | 0 | 1 |
| Archaea | Asgard | Candidatus Heimdallarchaeota | - | JQ817966 | 0 | 1 |
| Archaea | Asgard | Candidatus Heimdallarchaeota | - | DQ640139 | 0 | 1 |
| Archaea | Asgard | Candidatus Heimdallarchaeota | - | JX000774 | 0 | 1 |
| Archaea | Asgard | Candidatus Heimdallarchaeota | - | GU553642 | 0 | 1 |
| Archaea | Asgard | Candidatus Heimdallarchaeota | - | JQ989558 | 0 | 1 |
| Archaea | Asgard | Candidatus Lokiarchaeote | Prometheoarchaeum syntrophicum | CP042905 | 0 | 1 |
| Archaea | Asgard | Candidatus Lokiarchaeote | Lokiarchaeum sp. GC14_75 | JYIM01 | 1 | 1 |
| Archaea | Asgard | Candidatus Lokiarchaeote | Lokiarchaeota archaeon CR_4 | MBAA01 | 1 | 0 |
| Archaea | Asgard | Candidatus Lokiarchaeote | Lokiarchaeota archaeon Loki_b31 | NJBI01 | 1 | 0 |
| Archaea | Asgard | Candidatus Lokiarchaeote | Lokiarchaeota archaeon Loki_b32 | NJBH01 | 1 | 0 |
| Archaea | Asgard | Candidatus Lokiarchaeote | - | GU135483 | 0 | 1 |
| Archaea | Asgard | Candidatus Lokiarchaeote | - | JQ989629 | 0 | 1 |
| Archaea | Asgard | Candidatus Lokiarchaeote | - | EF125503 | 0 | 1 |
| Archaea | Asgard | Candidatus Lokiarchaeote | - | AB161349 | 0 | 1 |
| Archaea | Asgard | Candidatus Lokiarchaeote | - | AJ579317 | 0 | 1 |
| Archaea | Asgard | Candidatus Lokiarchaeote | - | HM244113 | 0 | 1 |
| Archaea | Asgard | Candidatus Lokiarchaeote | - | EU481570 | 0 | 1 |
| Archaea | Asgard | Candidatus Lokiarchaeote | - | AB301981 | 0 | 1 |
| Archaea | Asgard | Candidatus Lokiarchaeote | - | KX952655 | 0 | 1 |
| Archaea | Asgard | Candidatus Lokiarchaeote | - | KX952720 | 0 | 1 |
| Archaea | Asgard | Candidatus Lokiarchaeote | - | AY542580 | 0 | 1 |
| Archaea | Asgard | Candidatus Odinarchaeota | Odinarchaeota archaeon LCB_4 | MDVT01 | 1 | 1 |
| Archaea | Asgard | Candidatus Odinarchaeota | - | KX764885 | 0 | 1 |
| Archaea | Asgard | Candidatus Odinarchaeota | - | AB019719 | 0 | 1 |
| Archaea | Asgard | Candidatus Odinarchaeota | - | AY280451 | 0 | 1 |
| Archaea | Asgard | Candidatus Thorarchaeote | Thorarchaeota B41_G1 | QMYT01 | 0 | 1 |
| Archaea | Asgard | Candidatus Thorarchaeote | Thorarchaeota B29_G2 | QMYU01 | 0 | 1 |
| Archaea | Asgard | Candidatus Thorarchaeote | Thorarchaeota B59_G1 | QMYS01 | 0 | 1 |
| Archaea | Asgard | Candidatus Thorarchaeote | Thorarchaeota B65_G9 | QMYR01 | 0 | 1 |
| Archaea | Asgard | Candidatus Thorarchaeote | Thorarchaeote AB_25 | MEHG01 | 1 | 0 |
| Archaea | Asgard | Candidatus Thorarchaeote | Thorarchaeota archaeon SMTZ1 45 | LRSL01 | 1 | 1 |
| Archaea | Asgard | Candidatus Thorarchaeote | Thorarchaeota archaeon SMTZ1 83 | LRSK01 | 1 | 1 |
| Archaea | Asgard | Candidatus Thorarchaeote | Thorarchaeota archaeon MP11T_1 | PJET01 | 1 | 1 |
| Archaea | Asgard | Candidatus Thorarchaeote | Thorarchaeota archaeon MP8T_1 | PJER01 | 1 | 1 |
| Archaea | Asgard | Candidatus Thorarchaeote | Thorarchaeota archaeon MP9T_1 | PJES01 | 1 | 0 |
| Archaea | Asgard | Candidatus Thorarchaeote | Thorarchaeota archaeon SMTZ 45 | LRSM01 | 1 | 0 |
| Archaea | Asgard | Candidatus Thorarchaeote | - | BART01020584 | 0 | 1 |
| Archaea | Asgard | Candidatus Thorarchaeote | - | FJ351268 | 0 | 1 |
| Archaea | Asgard | Candidatus Thorarchaeote | - | FJ352077 | 0 | 1 |
| Archaea | Asgard | Candidatus Helarchaeota | Helarchaeota Hel_GB_B | SUPR01 | 1 | 0 |
| Archaea | Asgard | Candidatus Helarchaeota | Helarchaeota Hel_GB_A | SUPS01 | 1 | 0 |
| Archaea | Asgard | Asgard group archaeon | Gerdarchaeota_SZ_4_bin5_60 | GCA_011364945 | 1 | 0 |
| Archaea | Asgard | Asgard group archaeon | Gerdarchaeota_HMA_bin2_96 | GCA_013166835 | 1 | 0 |
| Archaea | Asgard | Asgard group archaeon | Gerdarchaeota_YT_2_057 | GCA_011366245 | 1 | 0 |
| Archaea | Asgard | Asgard group archaeon | Gerdarchaeota_MP5_2_1192 | GCA_011366295 | 1 | 0 |
| Archaea | Asgard | Asgard group archaeon | Gerdarchaeota_B18_G1 | GCA_003662935 | 1 | 0 |
| Archaea | Asgard | Asgard group archaeon | Gerdarchaeota_YT_5_010 | GCA_011366225 | 1 | 0 |
| Archaea | Asgard | Asgard group archaeon | Gerdarchaeota_YT_1_045 | GCA_011366285 | 1 | 0 |
| Archaea | DPANN | Candidatus Diapherotrites | Iainarchaeum andersonii SCGC AAA011 E11 | AQRS01 | 1 | 0 |
| Archaea | DPANN | Candidatus Micrarchaeota | Micrarchaeum acidiphilum ARMAN 2 | ACVJ01 | 1 | 0 |
| Archaea | DPANN | Nanoarchaeota | Nanoarchaeota archaeon SCGC AAA011 G17 | AQRL01 | 1 | 0 |
| Archaea | DPANN | Nanoarchaeota | Nanobsidianus stetteri | APJZ01 | 1 | 0 |
| Archaea | DPANN | Woesearchaeota | archaeon GW2011_AR15 | CP010425 | 1 | 0 |
| Archaea | DPANN | Woesearchaeota | archaeon GW2011_AR20 | CP010426 | 1 | 0 |
| Archaea | Euryarchaeota | Archaeoglobi | Archaeoglobus fulgidus DSM 4304 | AE000782 | 1 | 0 |
| Archaea | Euryarchaeota | Archaeoglobi | Euryarchaeota Ferroglobus placidus DSM 10642 | CP001899 | 1 | 0 |
| Archaea | Euryarchaeota | Methanobacteria | Methanobacterium sp. AL 21 | CP002551 | 1 | 0 |
| Archaea | Euryarchaeota | Methanobacteria | Methanosphaera stadtmanae DSM 3091 | CP000102 | 1 | 0 |
| Archaea | Euryarchaeota | Methanobacteria | Methanothermobacter thermautotrophicus str.Delta H | AE000666 | 1 | 0 |
| Archaea | Euryarchaeota | Methanomicrobia | Methanocella paludicola SANAE | AP011532 | 1 | 0 |
| Archaea | Euryarchaeota | Methanomicrobia | Methanoculleus marisnigri JR1 | CP000562 | 1 | 0 |
| Archaea | Euryarchaeota | Methanomicrobia | Methanolacinia petrolearia DSM 11571 | CP002117 | 1 | 0 |
| Archaea | Euryarchaeota | Methanomicrobia | Methanosarcina acetivorans C2A | AE010299 | 1 | 0 |
| Archaea | Euryarchaeota | Thermococci | Pyrococcus furiosus DSM 3638 | AE009950 | 1 | 1 |
| Archaea | Euryarchaeota | Thermococci | Thermococcus kodakarensis KOD1 | AP006878 | 1 | 1 |
| Archaea | Euryarchaeota | Thermoplasmata | Ferroplasma acidarmanus fer1 | CP004145 | 1 | 0 |
| Archaea | Euryarchaeota | Thermoplasmata | Methanomassiliicoccus luminyensis B10 | CAJE01 | 1 | 0 |
| Archaea | Euryarchaeota | Thermoplasmata | Thermoplasma acidophilum DSM 1728 | AL139299 | 1 | 1 |
| Archaea | Euryarchaeota | unclassified Euryarchaeota | Aciduliprofundum boonei T469 | CP001941 | 1 | 0 |
| Archaea | Euryarchaeota | uncultured marine group II euryarchaeote | uncultured marine group II euryarchaeote | CM001443 | 0 | 1 |
| Archaea | Euryarchaeota | Methanotorris | Methanotorris igneus Kol 5 | CP002737 | 0 | 1 |
| Archaea | Euryarchaeota | Picrophilus | Picrophilus torridus DSM 9790 | AE017261 | 0 | 1 |
| Archaea | TACK group | Korarchaeota | Candidatus Korarchaeum cryptofilum OPF8 | CP000968 | 1 | 0 |
| Archaea | TACK group | Korarchaeota | Korarchaeota | QNVG01 | 0 | 0 |
| Archaea | TACK group | Crenarchaeota | Ignicoccus hospitalis KIN4/I | CP000816 | 1 | 0 |
| Archaea | TACK group | Crenarchaeota | Aeropyrum pernix K1 | BA000002 | 1 | 0 |
| Archaea | TACK group | Crenarchaeota | Caldivirga maquilingensis IC 167 | CP000852 | 1 | 0 |
| Archaea | TACK group | Crenarchaeota | Desulfurococcus kamchatkensis | CP001140 | 1 | 0 |
| Archaea | TACK group | Crenarchaeota | Metallosphaera cuprina Ar 4 | CP002656 | 1 | 0 |
| Archaea | TACK group | Crenarchaeota | Pyrobaculum aerophilum str. IM2 | AE009441 | 1 | 0 |
| Archaea | TACK group | Crenarchaeota | Pyrolobus fumarii 1A | CP002838 | 1 | 0 |
| Archaea | TACK group | Crenarchaeota | Sulfolobus acidocaldarius DSM 639 | CP000077 | 1 | 0 |
| Archaea | TACK group | Crenarchaeota | Thermofilum pendens Hrk 5 | CP000505 | 1 | 0 |
| Archaea | TACK group | Crenarchaeota | Thermoproteus uzoniensis 768 20 | CP002590 | 1 | 0 |
| Archaea | TACK group | Crenarchaeota | Vulcanisaeta distributa DSM 14429 | CP002100 | 1 | 0 |
| Archaea | TACK group | Aigarchaeota | Aigarchaeota archaeon JGI 0000106 J15 | ASPF01 | 1 | 0 |
| Archaea | TACK group | Aigarchaeota | Aigarchaeota archaeon SCGC AAA471 G05 | ASLV01 | 1 | 0 |
| Archaea | TACK group | Aigarchaeota | Candidatus Caldiarchaeum subterraneum | BA000048 | 0 | 0 |
| Archaea | TACK group | Thaumarchaeota | Candidatus Nitrososphaera gargensis Ga9.2 | CP002408 | 1 | 0 |
| Archaea | TACK group | Thaumarchaeota | Cenarchaeum symbiosum A | DP000238 | 1 | 0 |
| Archaea | TACK group | Thaumarchaeota | Nitrosopumilus maritimus SCM1 | CP000866 | 1 | 0 |
| Archaea | TACK group | Thaumarchaeota | Uncultured archaeon clone ASS_A1 | CP009479 | 0 | 0 |
| Archaea | TACK group | Thaumarchaeota | Candidatus Nitrosoarchaeum limnia SFB1 | CM001158 | 0 | 0 |
| Archaea | TACK group | Barhyarchaeota | Barhyarchaeota das_tool.concoct.117 | SPBR01 | 1 | 0 |
| Archaea | TACK group | Barhyarchaeota | Bathyarchaeota archaeon BA1 | LIHJ01 | 1 | 0 |

**Supplementary Table 5.** List of 56 ribosomal proteins used for inference of phylogenomic trees in this study.

| **arCOG** | **COG** | **Description** |
| --- | --- | --- |
| arCOG04086 | COG1841 | Ribosomal protein L30/L7E |
| arCOG00779 | COG0200 | Ribosomal protein L15 |
| arCOG04087 | COG0098 | Ribosomal protein S5 |
| arCOG04154 | COG2007 | Ribosomal protein S8E |
| arCOG04088 | COG0256 | Ribosomal protein L18 |
| arCOG04372 | COG0080 | Ribosomal protein L11 |
| arCOG01751 | COG1358 | Ribosomal protein L7Ae or related RNA K-turn-binding protein |
| arCOG04090 | COG0097 | Ribosomal protein L6P/L9E |
| arCOG04242 | COG0102 | Ribosomal protein L13 |
| arCOG04245 | COG0052 | Ribosomal protein S2 |
| arCOG04243 | COG0103 | Ribosomal protein S9 |
| arCOG04089 | COG2147 | Ribosomal protein L19E |
| arCOG04091 | COG0096 | Ribosomal protein S8 |
| arCOG04093 | COG1471 | Ribosomal protein S4E |
| arCOG04094 | COG0198 | Ribosomal protein L24 |
| arCOG04095 | COG0093 | Ribosomal protein L14 |
| arCOG04289 | COG0081 | Ribosomal protein L1 |
| arCOG00780 | COG1727 | Ribosomal protein L18E |
| arCOG00781 | COG1717 | Ribosomal protein L32E |
| arCOG04067 | COG0090 | Ribosomal protein L2 |
| arCOG04070 | COG0087 | Ribosomal protein L3 |
| arCOG04071 | COG0088 | Ribosomal protein L4 |
| arCOG04092 | COG0094 | Ribosomal protein L5 |
| arCOG04096 | COG0186 | Ribosomal protein S17 |
| arCOG04288 | COG0244 | Ribosomal protein L10 |
| arCOG04072 | COG0089 | Ribosomal protein L23 |
| arCOG04097 | COG0092 | Ribosomal protein S3 |
| arCOG04098 | COG0091 | Ribosomal protein L22 |
| arCOG04239 | COG0522 | Ribosomal protein S4 or related protein |
| arCOG04314 | COG2053 | Ribosomal protein S28E/S33 |
| arCOG04182 | COG2004 | Ribosomal protein S24E |
| arCOG01946 | COG2125 | Ribosomal protein S6E (S10) |
| arCOG04129 | COG2139 | Ribosomal protein L21E |
| arCOG04186 | COG1890 | Ribosomal protein S3AE |
| arCOG04255 | COG0048 | Ribosomal protein S12 |
| arCOG00785 | COG0255 | Ribosomal protein L29 |
| arCOG01752 | COG1841 | Ribosomal protein L30/L7E |
| arCOG04208 | COG1997 | Ribosomal protein L37AE/L43A |
| arCOG01758 | COG0051 | Ribosomal protein S10 |
| arCOG04113 | COG0197 | Ribosomal protein L16/L10AE |
| arCOG04240 | COG0100 | Ribosomal protein S11 |
| arCOG04254 | COG0049 | Ribosomal protein S7 |
| arCOG04287 | COG2058 | Ribosomal protein L12E/L44/L45/RPP1/RPP2 |
| arCOG04099 | COG0185 | Ribosomal protein S19 |
| arCOG04185 | COG0184 | Ribosomal protein S15P/S13E |
| arCOG01722 | COG0099 | Ribosomal protein S13 |
| arCOG04209 | COG1632 | Ribosomal protein L15E |
| arCOG04473 | COG2097 | Ribosomal protein L31E |
| arCOG01885 | COG1383 | Ribosomal protein S17E |
| arCOG01344 | COG2238 | Ribosomal protein S19E (S16A) |
| arCOG04108 | COG2051 | Ribosomal protein S27E |
| arCOG04109 | COG1631 | Ribosomal protein L44E |
| arCOG04167 | COG0093 | Ribosomal protein L14 |
| arCOG04183 | COG1998 | Ribosomal protein S27AE |
| arCOG01950 | COG2075 | Ribosomal protein L24E |
| arCOG04175 | COG2157 | Ribosomal protein L20A (L18A) |

**Supplementary Table 10.** Relative abundance of Hermodarchaeota bins in sequencing data.

| **Bins** | **Samples** | **Number of raw_reads** | **Number of mapped_reads** | **Relative abundance**  **(%)** |
| --- | --- | --- | --- | --- |
| h02s_80 | h02s | 201686883 | 337935 | 0.168% |
| h02s_80 | h02m | 223296758 | 508505 | 0.228% |
| h02s_80 | h02b | 204437974 | 122585 | 0.060% |
| h02s_80 | h03s | 222631731 | 103453 | 0.046% |
| h02s_80 | h03m | 244015268 | 49273 | 0.020% |
| h02s_80 | h03b | 227434540 | 34334 | 0.015% |
| h02s_68 | h02s | 201686883 | 425141 | 0.211% |
| h02s_68 | h02m | 223296758 | 839717 | 0.376% |
| h02s_68 | h02b | 204437974 | 960500 | 0.470% |
| h02s_68 | h03s | 222631731 | 438965 | 0.197% |
| h02s_68 | h03m | 244015268 | 538602 | 0.221% |
| h02s_68 | h03b | 227434540 | 259460 | 0.114% |
| h02m_131 | h02s | 201686883 | 271708 | 0.135% |
| h02m_131 | h02m | 223296758 | 705112 | 0.316% |
| h02m_131 | h02b | 204437974 | 984284 | 0.481% |
| h02m_131 | h03s | 222631731 | 382833 | 0.172% |
| h02m_131 | h03m | 244015268 | 498596 | 0.204% |
| h02m_131 | h03b | 227434540 | 315804 | 0.139% |

**Supplementary Table 11.** Blastn output results retrieved by comparing Hermodarchaeota 16S rRNA gene sequence with those from sediment samples and published Asgard archaea.

| **Query id** | **Subject id** | **% identity** | **alignment length** | **mismatches** | **gap openings** | **q. start** | **q. end** | **s. start** | **s. end** | **evalue** | **bit score** |
| --- | --- | --- | --- | --- | --- | --- | --- | --- | --- | --- | --- |
| h02s_26_NODE_7530 | h02s_matam_3940 | 95.775 | 1065 | 43 | 2 | 3 | 1066 | 1 | 1064 | 0 | 1716 |
| h02s_26_NODE_7530 | h02s_matam_8111 | 95.108 | 879 | 43 | 0 | 69 | 947 | 1 | 879 | 0 | 1386 |
| h02s_26_NODE_7530 | Odinarchaeote-LCB_4_MDVT01000007.1 | 83.711 | 927 | 141 | 10 | 145 | 1066 | 186 | 1107 | 0 | 867 |
| h02s_26_NODE_7530 | Lokiarchaeota-archaeon-B53_G9SDNY01000025.1 | 79.835 | 848 | 152 | 18 | 227 | 1066 | 219 | 1055 | 1.46E-174 | 601 |
| h02s_26_NODE_7530 | Lokiarchaeum-sp.-GC14_75_JYIM01000321.1 | 79.762 | 756 | 129 | 22 | 321 | 1066 | 1 | 742 | 2.52E-152 | 527 |
| h02s_26_NODE_7530 | Thorarchaeota-archaeon-MP11T_1_PJET01000033.1 | 77.39 | 774 | 150 | 20 | 286 | 1046 | 281 | 1042 | 4.36E-125 | 436 |
| h02s_26_NODE_7530 | Thorarchaeota-archaeon-MP8T_1_PJER01000019.1 | 77.261 | 774 | 151 | 22 | 286 | 1046 | 281 | 1042 | 2.03E-123 | 431 |
| h02s_26_NODE_7530 | Helarchaeota_Ga0180301_100789461 | 81.683 | 404 | 64 | 10 | 480 | 878 | 1 | 399 | 2.74E-92 | 327 |
| h02s_26_NODE_7530 | Heimdalarchaeote-LC_3_MDVS01000157.1 | 72.931 | 713 | 162 | 28 | 369 | 1066 | 6 | 702 | 1.72E-59 | 219 |
| h02m_131_k127_628877 | Odinarchaeote-LCB_4_MDVT01000007.1 | 79.921 | 508 | 72 | 21 | 1 | 505 | 1063 | 1543 | 3.53E-98 | 346 |
| h02m_131_k127_628877 | Helarchaeota_Ga0180301_100789461 | 77.559 | 508 | 84 | 17 | 1 | 505 | 577 | 1057 | 3.63E-78 | 279 |
| h02m_131_k127_628877 | Lokiarchaeum-sp.-GC14_75_JYIM01000321.1 | 78.61 | 374 | 75 | 4 | 1 | 372 | 698 | 1068 | 4.77E-67 | 243 |
| h02m_131_k127_628877 | Lokiarchaeota-archaeon-B53_G9SDNY01000025.1 | 77.807 | 374 | 78 | 4 | 1 | 372 | 1011 | 1381 | 4.8E-62 | 226 |

**Supplementary Table 14.** Comparison of alkylsuccinate synthase (Ass)/benzylsuccinate synthase (Bss) of Hermodarchaeota with known Ass/Bss and pyruvate formate lyase (pfl) using Blastp.

| **Query id** | **Subject id** | **% identity** | **alignment length** | **mismatches** | **gap openings** | **q. start** | **q. end** | **s. start** | **s. end** | **evalue** | **bit score** |
| --- | --- | --- | --- | --- | --- | --- | --- | --- | --- | --- | --- |
| h02s_68 k137_1286063_3 | AssA1 from *D. alkenivorans* strain AK-01(ABH11460) | 33.38 | 605 | 380 | 12 | 52 | 642 | 235 | 830 | 1.18E-98 | 312 |
| h02s_68 k137_1286063_3 | BssA from *A. aromaticum* EbN1 (YP_158060) | 31.72 | 618 | 384 | 16 | 52 | 643 | 241 | 846 | 3.93E-87 | 281 |
| h02s_68 k137_1286063_3 | pfl from *E. coli* (NP_415423) | 25.08 | 650 | 414 | 20 | 10 | 646 | 169 | 758 | 3.94E-46 | 165 |
| h02s_68 k137_1286063_3 | pflD from *Archaeoglobus fulgidus* (AAB89800) | 38.15 | 637 | 383 | 8 | 144 | 771 | 7 | 641 | 8.74e-158 | 463 |
| h02m_131 k127_726590_1 | AssA1 from *D. alkenivorans* strain AK-01(ABH11460) | 29.22 | 705 | 447 | 14 | 128 | 794 | 139 | 829 | 7.03E-79 | 262 |
| h02m_131 k127_726590_1 | BssA from *A. aromaticum* EbN1 (YP_158060) | 30.65 | 620 | 373 | 18 | 209 | 794 | 248 | 844 | 1.03E-68 | 234 |
| h02m.131 k127_726590_1 | pfl from *E. coli* (NP_415423) | 27.29 | 590 | 354 | 21 | 229 | 794 | 216 | 754 | 2.21E-38 | 143 |
| h02m_131 k127_726590_1 | pflD from *Archaeoglobus fulgidus* (AAB89800) | 33.44 | 646 | 383 | 16 | 155 | 773 | 171 | 796 | 3.66e-107 | 336 |
| h02s_68 k137_157306_13 | AssA1 from *D. alkenivorans* strain AK-01(ABH11460) | 30.51 | 790 | 505 | 19 | 27 | 789 | 60 | 832 | 2.28E-96 | 310 |
| h02s_68 k137_157306_13 | BssA from *A. aromaticum* EbN1 (YP_158060) | 29.96 | 801 | 502 | 23 | 27 | 790 | 70 | 848 | 3.24E-90 | 294 |
| h02s_68 k137_157306_13 | pfl from *E. coli* (NP_415423) | 24.55 | 554 | 356 | 16 | 257 | 792 | 251 | 760 | 3.49E-29 | 114 |
| h02s_68 k137_157306_13 | pflD from *Archaeoglobus fulgidus* (AAB89800) | 33.80 | 787 | 498 | 17 | 4 | 773 | 8 | 788 | 8.06e-140 | 421 |

**Supplementary Table 15.** Comparison of alkylsuccinate synthase/benzylsuccinate synthase activating enzyme (Ass/Bss AE) of Hermodarchaeota with known Ass/Bss and pyruvate formate lyase (pfl) AE using Blastp.

| **Query id** | **Subject id** | **% identity** | **alignment length** | **mismatches** | **gap openings** | **q. start** | **q. end** | **s. start** | **s. end** | **evalue** | **bit score** |
| --- | --- | --- | --- | --- | --- | --- | --- | --- | --- | --- | --- |
| h02s.80 k137_2455425_12 | AssD2 from *D. alkenivorans* strain AK-01 (YP_002431363) | 39.16 | 286 | 160 | 3 | 1 | 276 | 19 | 300 | 1.30E-67 | 203 |
| h02s.80 k137_2455425_12 | AssD2' from *D. alkenivorans* strain AK-01 (YP_002429341) | 40.94 | 298 | 167 | 4 | 1 | 289 | 17 | 314 | 3.44E-86 | 250 |
| h02s.80 k137_2455425_12 | PflC from *Archaeoglobus fulgidus* (KUJ94427) | 40.48 | 252 | 146 | 2 | 1 | 252 | 20 | 267 | 3.56E-68 | 204 |
| h02s.80 k137_2455425_12 | BssD from *T. aromatica* K172 (CAA05050) | 38.46 | 260 | 152 | 4 | 1 | 253 | 18 | 276 | 3.47E-65 | 197 |
| h02s.80 k137_2455425_12 | pflD from *E.coli* (NP_415422) | 31.64 | 177 | 118 | 2 | 86 | 260 | 48 | 223 | 1.08E-31 | 107 |
| h02s.68 k137_3621160_2 | AssD2 from *D. alkenivorans* strain AK-01 (YP_002431363) | 42.91 | 282 | 151 | 3 | 23 | 294 | 6 | 287 | 9.42E-75 | 223 |
| h02s.68 k137_3621160_2 | AssD2' from *D. alkenivorans* strain AK-01 (YP_002429341) | 44.87 | 312 | 163 | 4 | 22 | 324 | 3 | 314 | 8.71E-98 | 281 |
| h02s.68 k137_3621160_2 | PflC from *Archaeoglobus fulgidus* (KUJ94427) | 41.26 | 269 | 154 | 2 | 19 | 287 | 3 | 267 | 1.54E-72 | 216 |
| h02s.68 k137_3621160_2 | BssD from *T. aromatica* K172 (CAA05050) | 37.59 | 274 | 165 | 2 | 20 | 287 | 2 | 275 | 1.31E-68 | 207 |
| h02s.68 k137_3621160_2 | pflD from *E.coli* (NP_415422) | 31.82 | 264 | 121 | 4 | 21 | 282 | 4 | 210 | 3.92E-40 | 131 |
| h02m.131 k127_726590_2 | AssD2 from *D. alkenivorans* strain AK-01 (YP_002431363) | 39.37 | 287 | 164 | 1 | 7 | 283 | 2 | 288 | 1.60E-73 | 218 |
| h02m.131 k127_726590_2 | AssD2' from *D. alkenivorans* strain AK-01 (YP_002429341) | 45.33 | 300 | 157 | 3 | 10 | 302 | 3 | 302 | 7.74E-100 | 285 |
| h02m.131 k127_726590_2 | PflC from *Archaeoglobus fulgidus* (KUJ94427) | 40.23 | 266 | 155 | 2 | 10 | 275 | 6 | 267 | 2.08E-72 | 215 |
| h02m.131 k127_726590_2 | BssD from *T. aromatica* K172 (CAA05050) | 38.38 | 271 | 161 | 3 | 12 | 276 | 6 | 276 | 9.81E-68 | 204 |
| h02m.131 k127_726590_2 | pflD from *E.coli* (NP_415422) | 30.42 | 263 | 124 | 5 | 10 | 270 | 5 | 210 | 3.86E-37 | 122 |

**Legends for Suppl. Tables 6-9, 12-13 and 16.**

**Please note that these Suppl. Tables are provided in separate excel files. For Suppl. Tables 6-8, they are placed into a excel file referred to as “Supplementary Tables_6-8.xlsx”**

**Supplementary Table 6.** Transcripts identified in metatranscriptomic datasets from mangrove wetlands when genes involved in important metabolic processes in the bin h02s_68 were used as reference sequences.

**Supplementary Table 7.** Transcripts identified in metatranscriptomic datasets from mangrove wetlands when genes involved in important metabolic processes in the bin h02s_80 were used as reference sequences.

**Supplementary Table 8.** Transcripts identified in metatranscriptomic datasets from mangrove wetlands when genes involved in important metabolic processes in the bin h02m_131 were used as reference sequences.

**Supplementary Table 9.** Metagenomic samples from a variety of environments containing homologues of Hermodarchaeota alkyl/benzyl-succinate synthase (Ass/Bss) and/or 16S rRNA genes.

**Supplementary Table 12.** Distribution of eukaryotic signature proteins in Hermodarchaeota and other Asgard archaea.

**Supplementary Table 13.** The genes used for metabolic reconstruction in this study.

**Supplementary Table 16.** Features of homologs of Hermodarchaeota alkyl/benzyl-succinate synthase (Ass/Bss) identified using Blasp in metagenomes from the six sediment samples from mangrove swamps in Techeng Island, China.

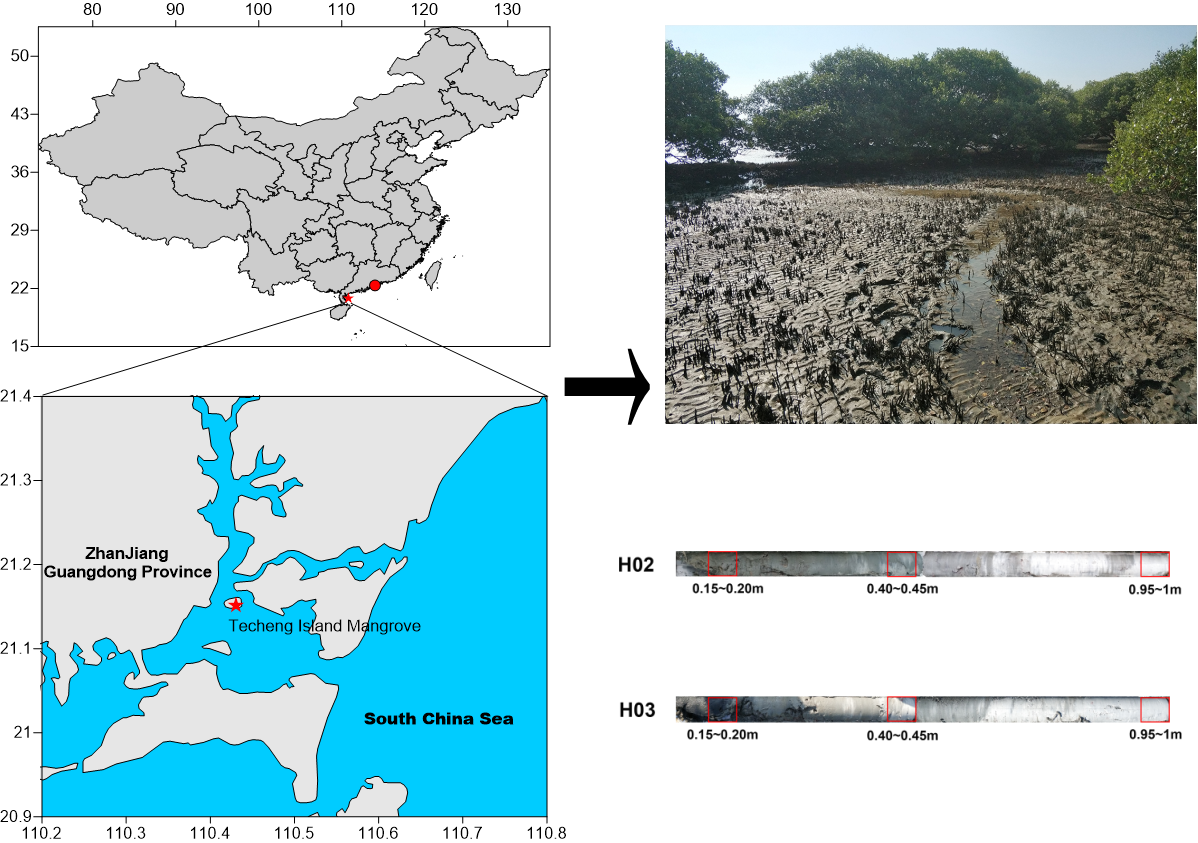

**Supplementary Fig. 1** Geographic locations of the sampling sites from mangrove swamps. The red stars represent the sites for metagenomic data in this study. The red dot represents the sites for metatranscriptomic data which are downloaded from NCBI database [33].

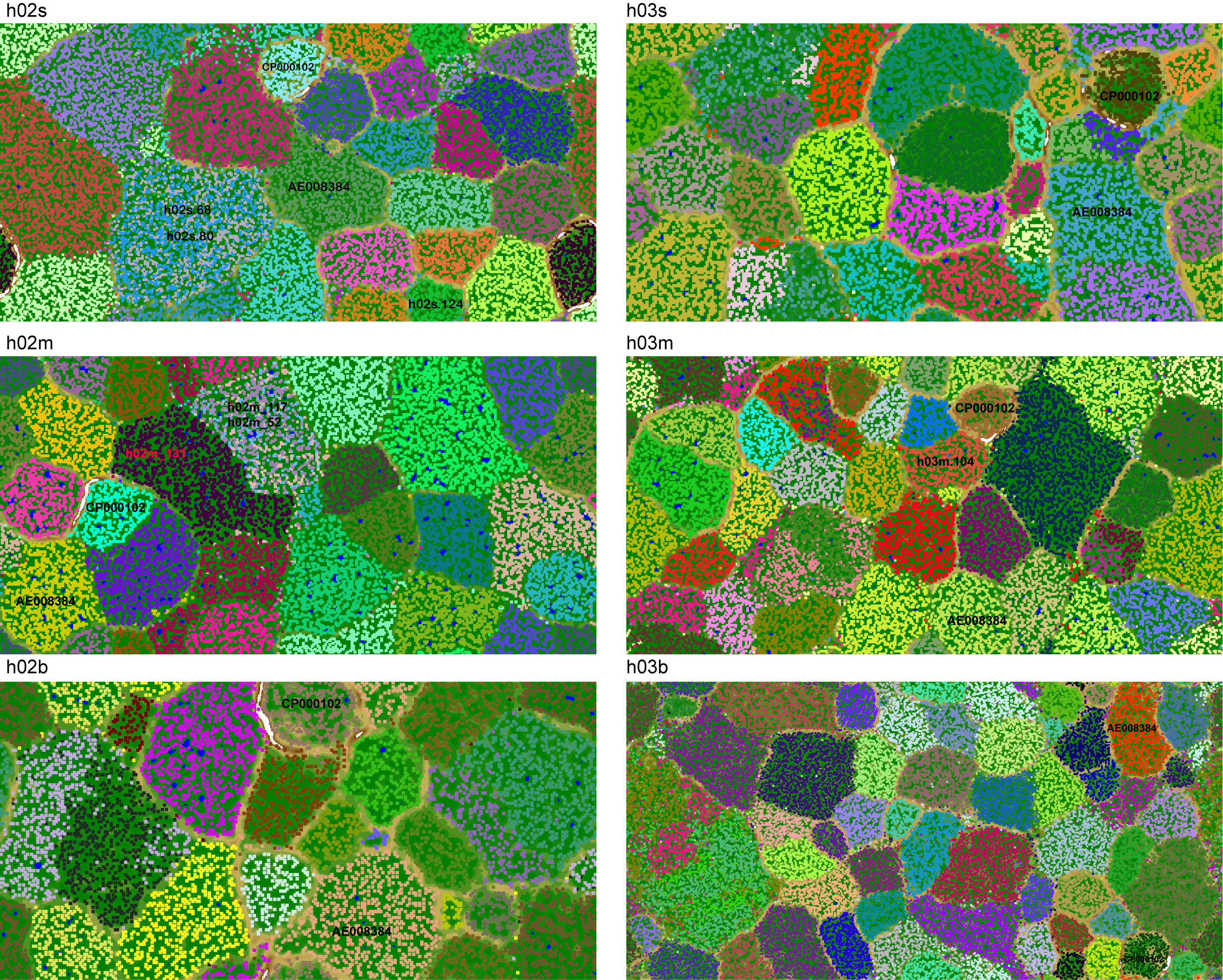

**Supplementary Fig. 2** Visualization of bins assembled from six mangrove sediment samples using Emergent Self-Organizing map (ESOM). Seven Hermodarchaeota bins were indicated, including h02s.80, h02m.131, h02s.68, h02s.124, h02m.52, h02m.117 and h03m_104. AE008384 and CP000102 represent genomes of *Methanosarchina mazei* strain Goe1 [34] and *Methanosphaera stadtmanae* [35]*,* respectively, and they were used as reference genomes.

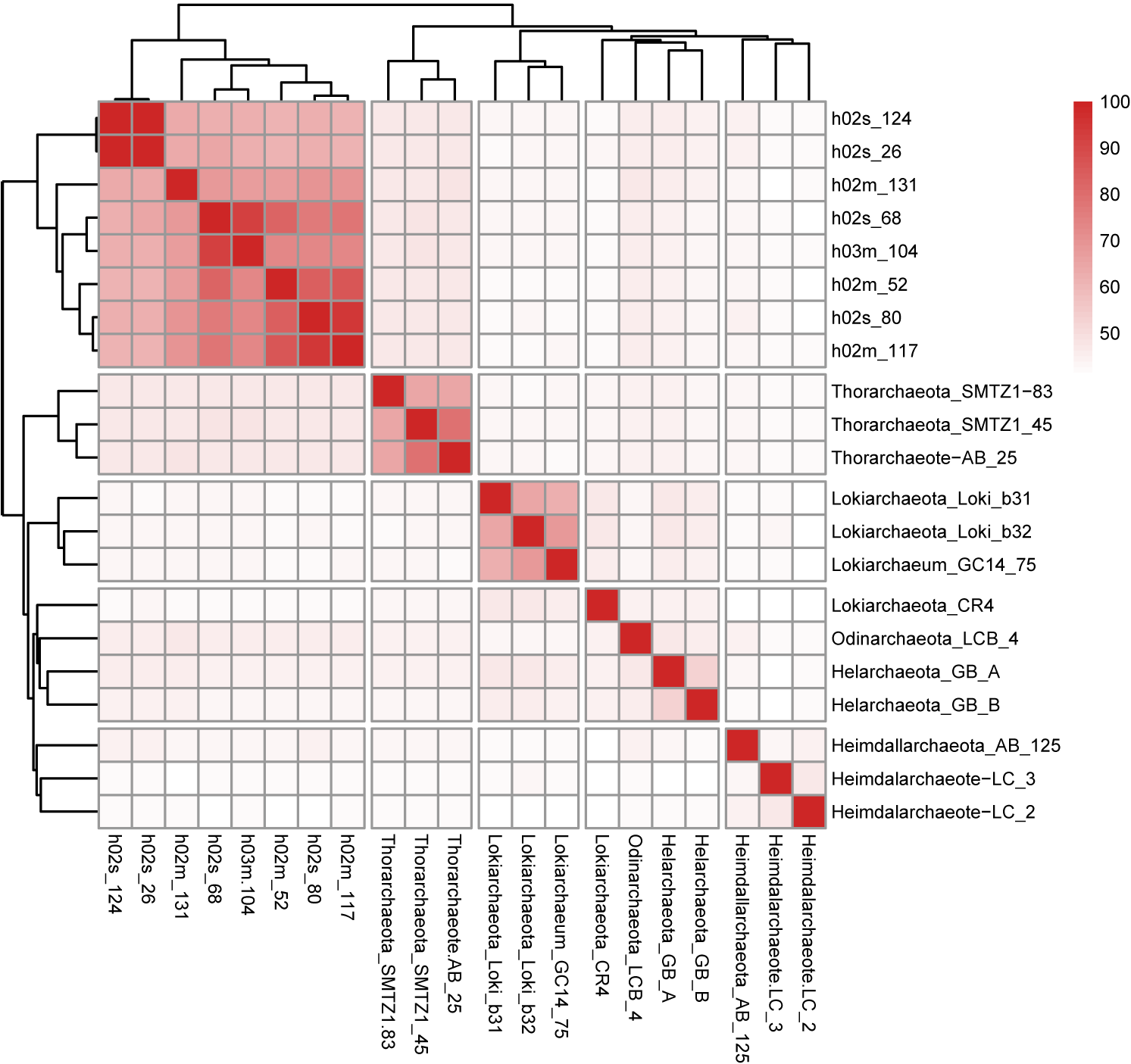

**Supplementary Fig. 3** Comparison of average amino acid identity (AAI) between Hermodarchaeota bins and published Asgard genomes. AAI was analyzed by CompareM.

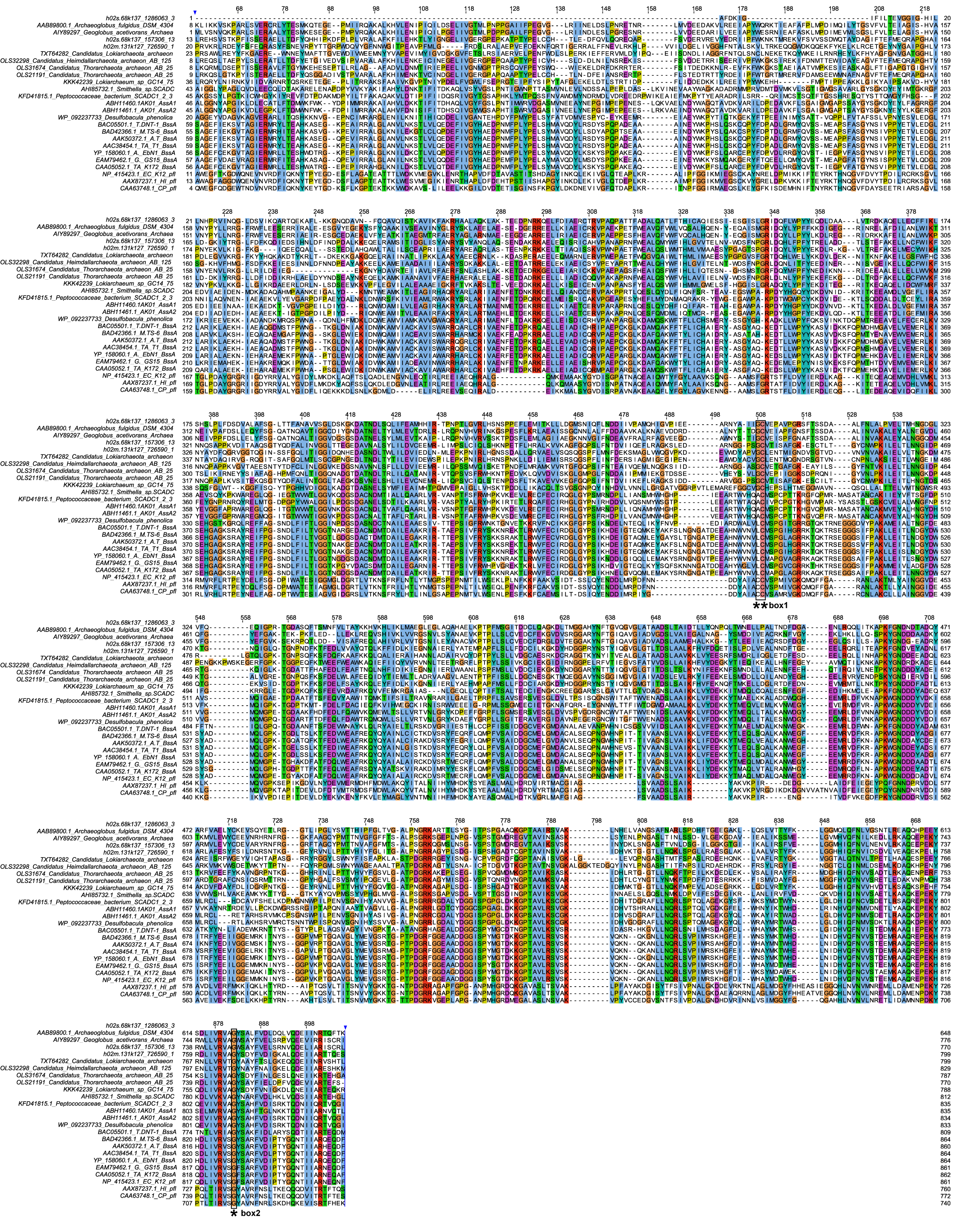

**Supplementary Fig. 4** Multiple sequence alignment of alkylsuccinate synthase (Ass)/benzylsuccinate synthase (Bss) of Hermodarchaeota (h02s_68 k137_1286063_3, h02s_68 k137_157306_13, h02m_131 k127_726590_1) with known Ass/Bss and pyruvate formate lyase (pfl). Uninterrupted sequences are shown except for the N-terminal and C-terminal variable regions. Conserved residues are shaded in colors. Box1 and box2 correspond to the conserved cysteine residues accepting the radical from the glycyl residue [36] and the conserved glycine residue, respectively. D. A_ AK-01, *D. alkenivorans* strain AK-01.

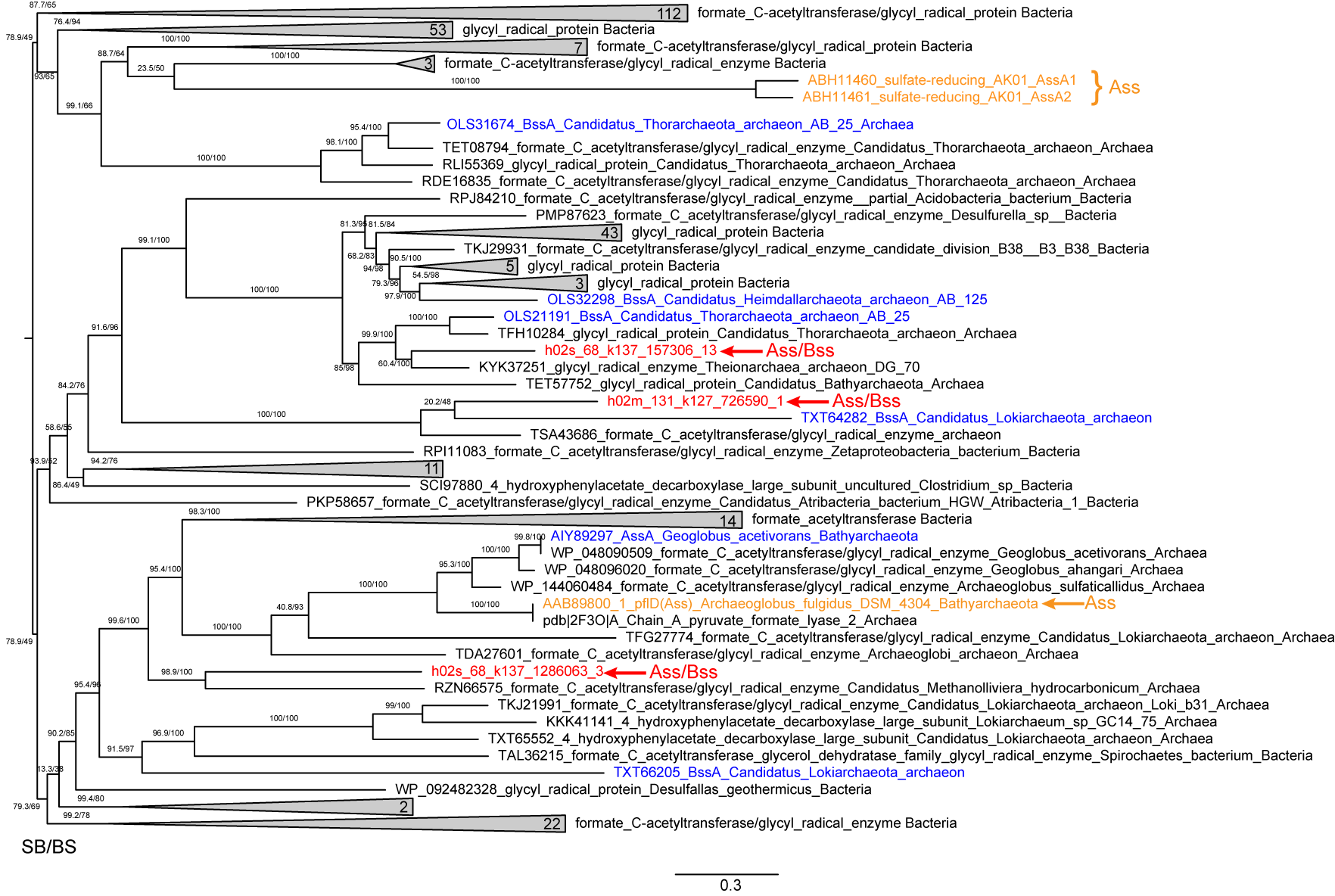

**Supplementary Fig. 5** Maximum-likelihood tree of alkyl/benzyl-succinate synthases (Ass/Bss) identified in Hermodarchaeota genomes and homologues from nr database reconstructed using IQtree with LG+G4 substitution model. Hermodarchaeota alkyl/benzyl-succinate synthases were red-coded. The verified alkyl-succinate synthases were shaded in yellow. The potential alkyl-succinate synthases were shaded in blue. The SH-like approximate likelihood ratio test (SB) and ultrafast bootstrap (BS) support values were shown.

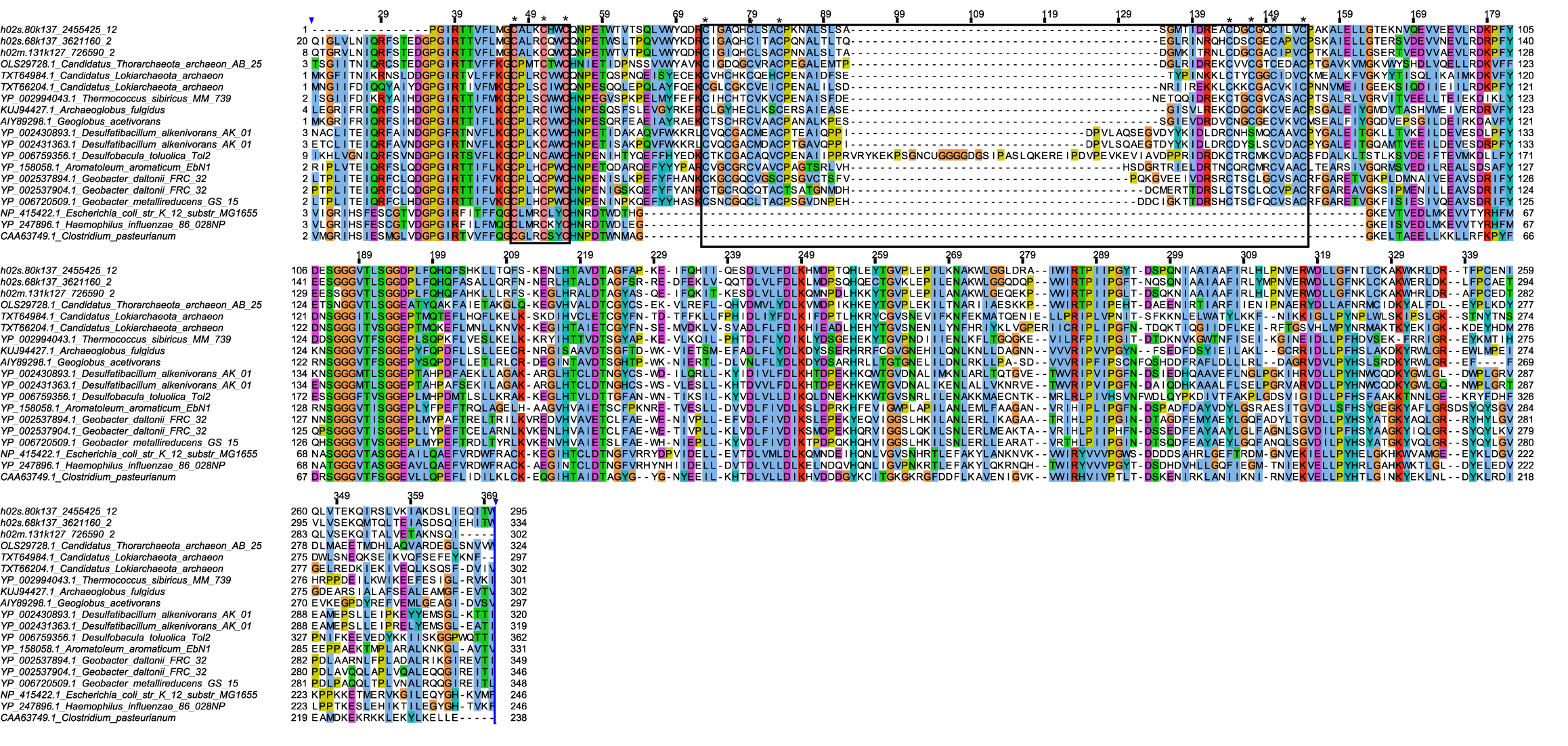

**Supplementary Fig. 6** Multiple sequence alignment of Hermodarchaeota Ass or Bss-activating enzyme (Ass/Bss AE) with known Ass/Bss and pyruvate formate lyase (pfl) AE. Uninterrupted sequences are shown except for the N-terminal and C-terminal variable regions. Boxes 1 and 2 correspond to the CxxxCxxC sequence motif and two cysteine-rich regions, respectively, and they are involved FeS cluster binding. Conserved residues are shaded in colors.

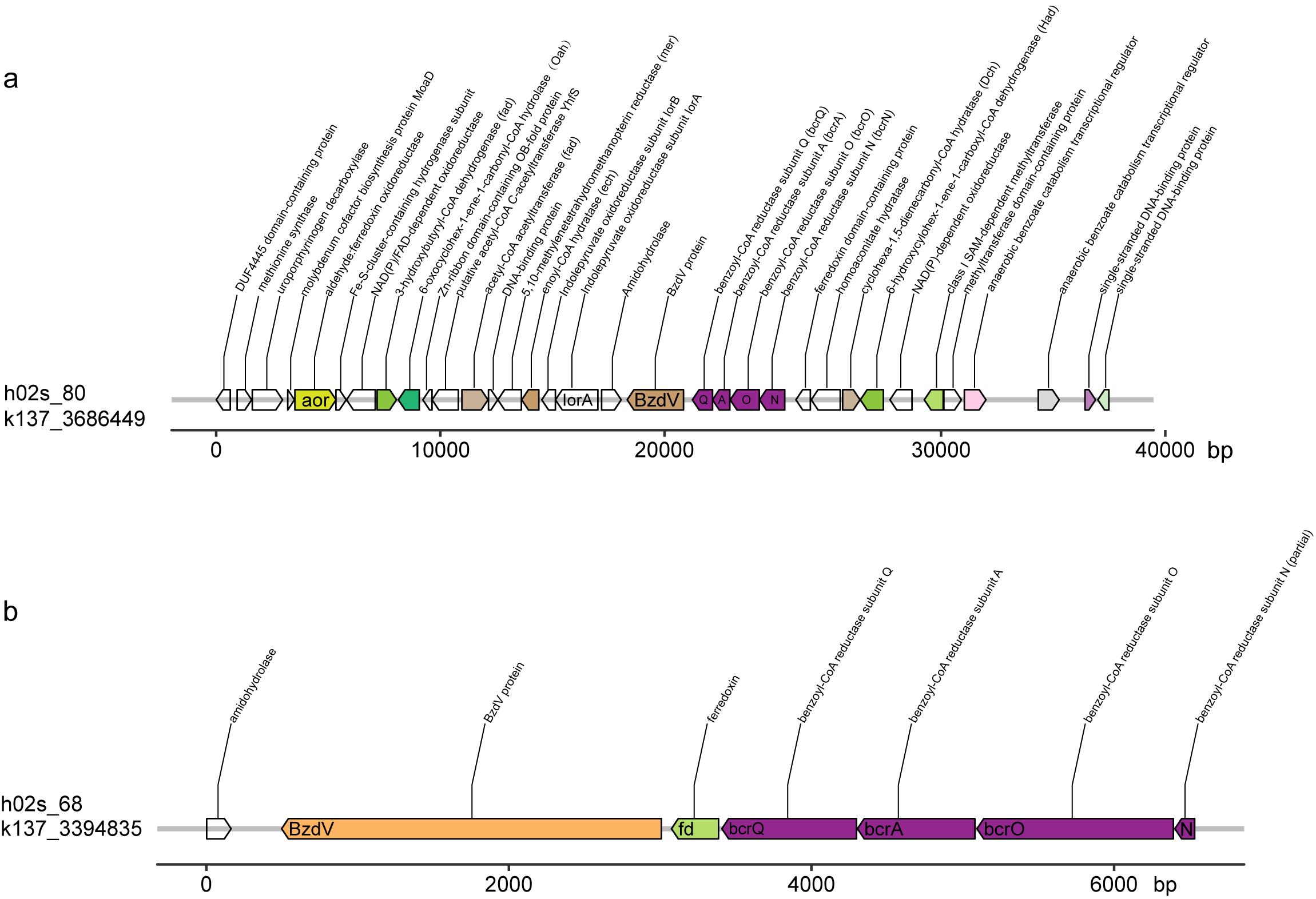

**Supplementary Fig. 7** The gene composition in the contigs containing the benzoyl-CoA reductase operon in h02s_80 and h02s_68 bins. Purple bocks indicate four subunits of the benzoyl-CoA reductase. Arrows show transcriptional orientation of the genes.

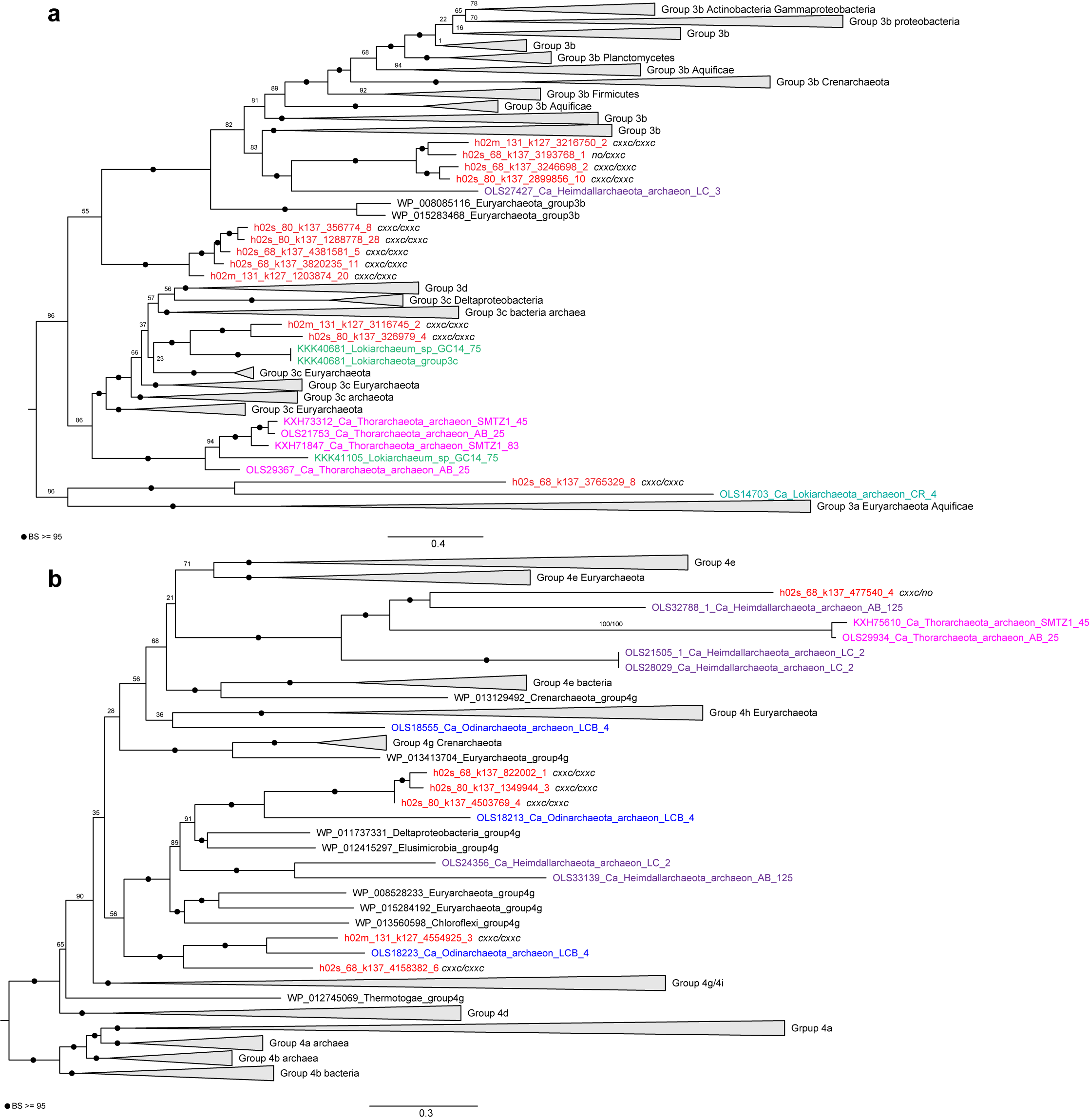

**Supplementary Fig. 8** Maximum-likelihood trees of the large subunit of group 3 and group 4 [NiFe]-hydrogenases reconstructed using IQtree v.1.6.12 with the best-fit model. **a** the large subunit of group 3 [NiFe]-hydrogenase (>300 amino acids); **b** the large subunit of group 4 [NiFe]-hydrogenase (>300 amino acids). Asgard hydrogenases were shaded in color as follows: Hermodarchaeota in red, Heimdallarchaeota in purple, Lokiarchaeota in green, Thorarchaetota in pink, and Odinarchaeum in blue. The N-terminal and C-terminal CxxC motifs were shown (two motifs: CxxC/ CxxC, one motif: no/ CxxC or CxxC/ no. The numbers at the nodes indicate ultrafast bootstrap values (BS). The ultrafast bootstrap values ≥ 95 are shown with black circles.

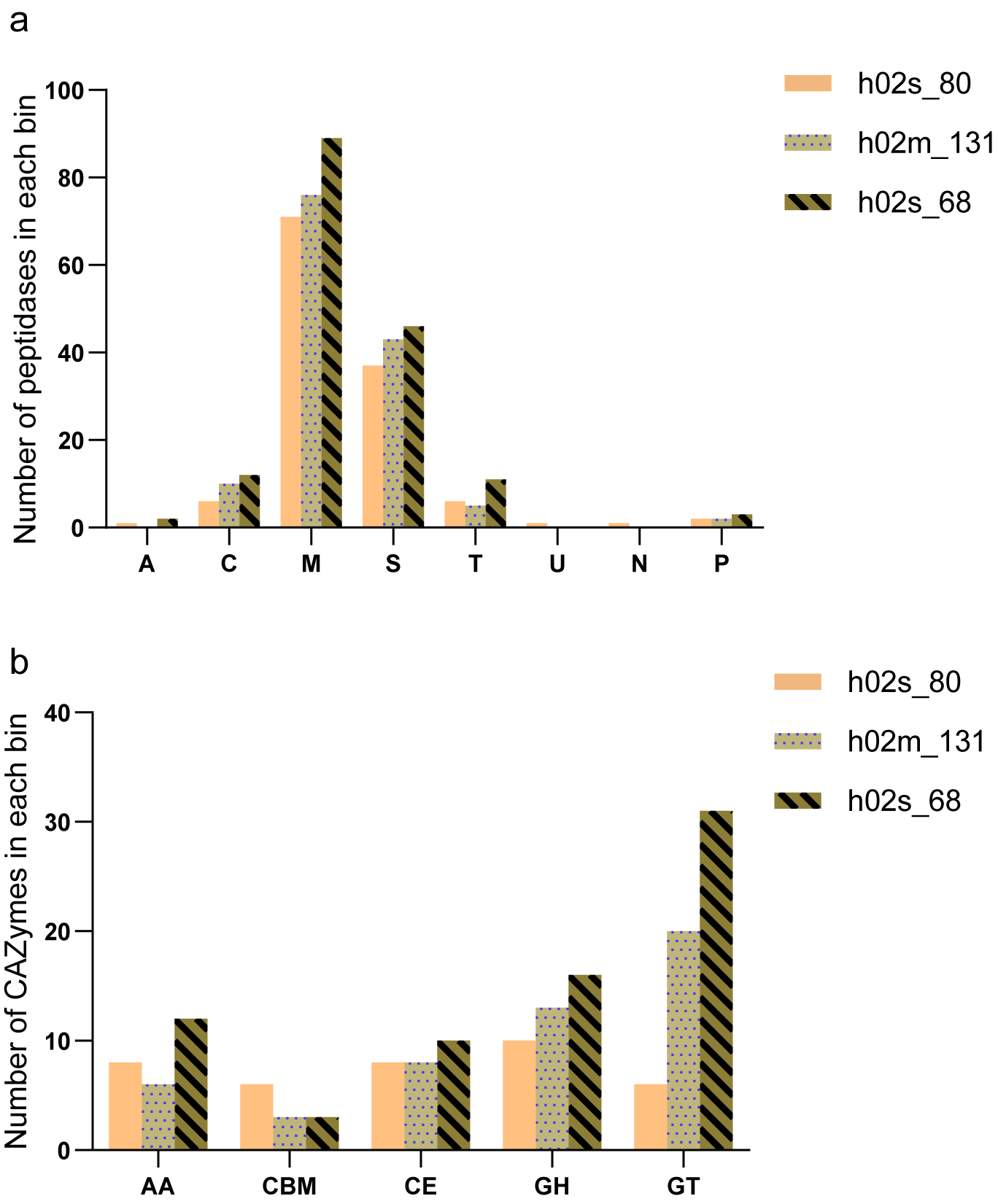

**Supplementary Fig. 9** Number of peptidases (a) and carbohydrate-active enzymes (CAZymes) (b) in three Hermodarchaeota genomes. Peptidases and CAZymes were identified using MEROPS database and dbCAN web server using default parameters, respectively. A aspartic peptidase; C, cysteine peptidase; M, metallopeptidase; S, serine peptidase; T, threonine peptidase; U, unknown catalytic type; N, asparagine peptide lyase; P, mixed peptidase. AA, auxiliary activity; CBM, carbohydrate-binding module; CE, carbohydrate esterase; GH, glycoside hydrolase; GT, glycosyltransferase.

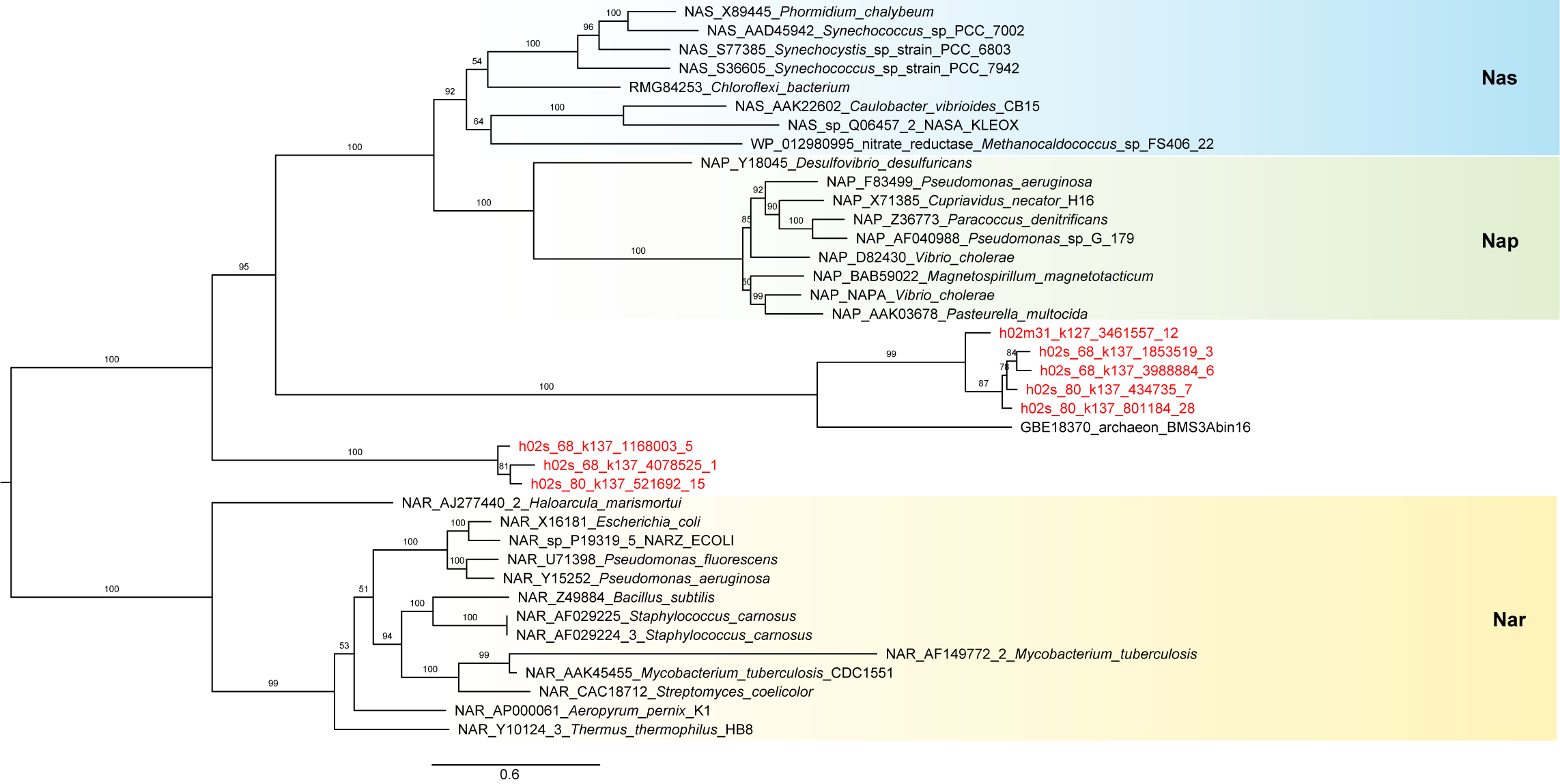

**Supplementary Fig. 10** Phylogenetic placement of nitrate reductases identified in Hermodarchaeota genomes. Maximum-likelihood tree was reconstructed using IQtree with under LG+ I+ G4 substitution model. A set of homologues of Nar, Nap and Nas from representative prokaryotic organisms were derived from a previous study [30]. Nitrate reductases from Hermodarchaeota were red-coded. Nas, prokaryotic assimilatory nitrate reductase; Nap, the periplasmic nitrate reductase; Nar, the membrane-associated prokaryotic nitrate reductase. The ultrafast bootstrap support values are shown.

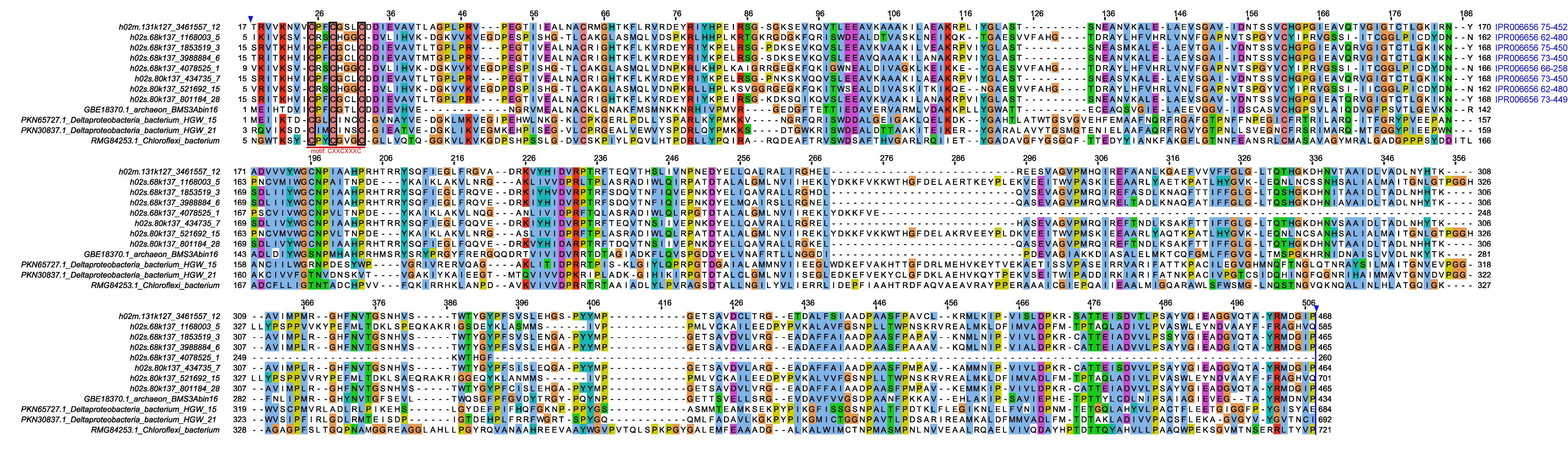

**Supplementary Fig. 11** Multiple sequence alignment of Hermodarchaeota nitrate reductases and homologues from nr database. These sequences contained a molybdopterin oxidoreductase domain (IPR006656). The CXXCXXXC motif which is required for all periplasmic nitrate reductase (Nap) is shown. Conserved residues are shaded in colors.

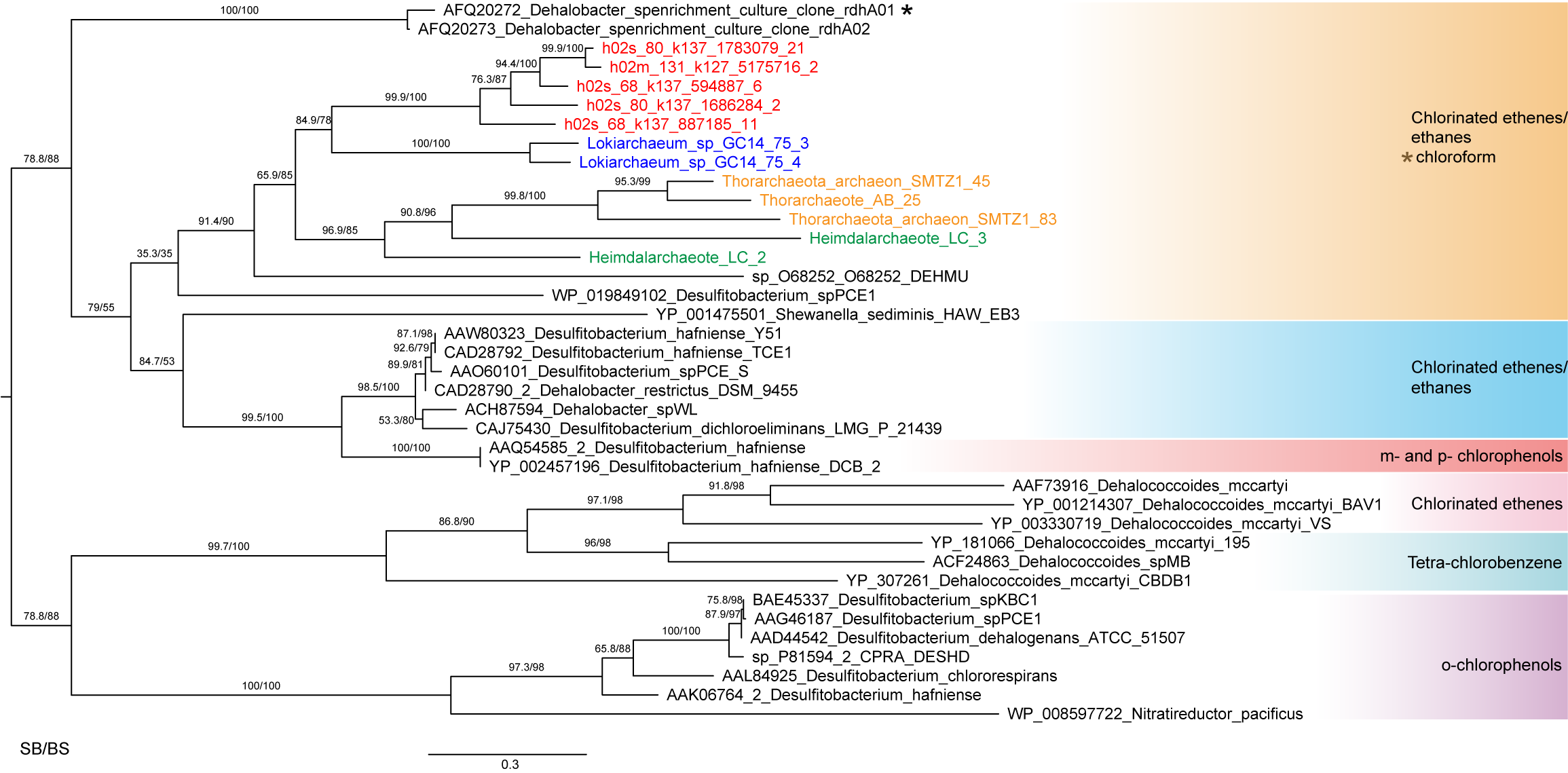

**Supplementary Fig. 12** Phylogenetic placement of reductive dehalogenases identified in Hermodarchaeota genomes. Functionally characterized reductive dehalogenases from bacteria were derived from a previous study [29]. Maximum-likelihood tree was reconstructed using IQtree with LG+I+G4 model. Asgard reductive dehalogenases were shaded in color as follows: Hermodarchaeota in red, Lokiarchaeota in blue, Thorarchaetota in yellow, Heimdallarchaeota in green. The SH-like approximate likelihood ratio test (SB) and ultrafast bootstrap (BS) support values were shown.

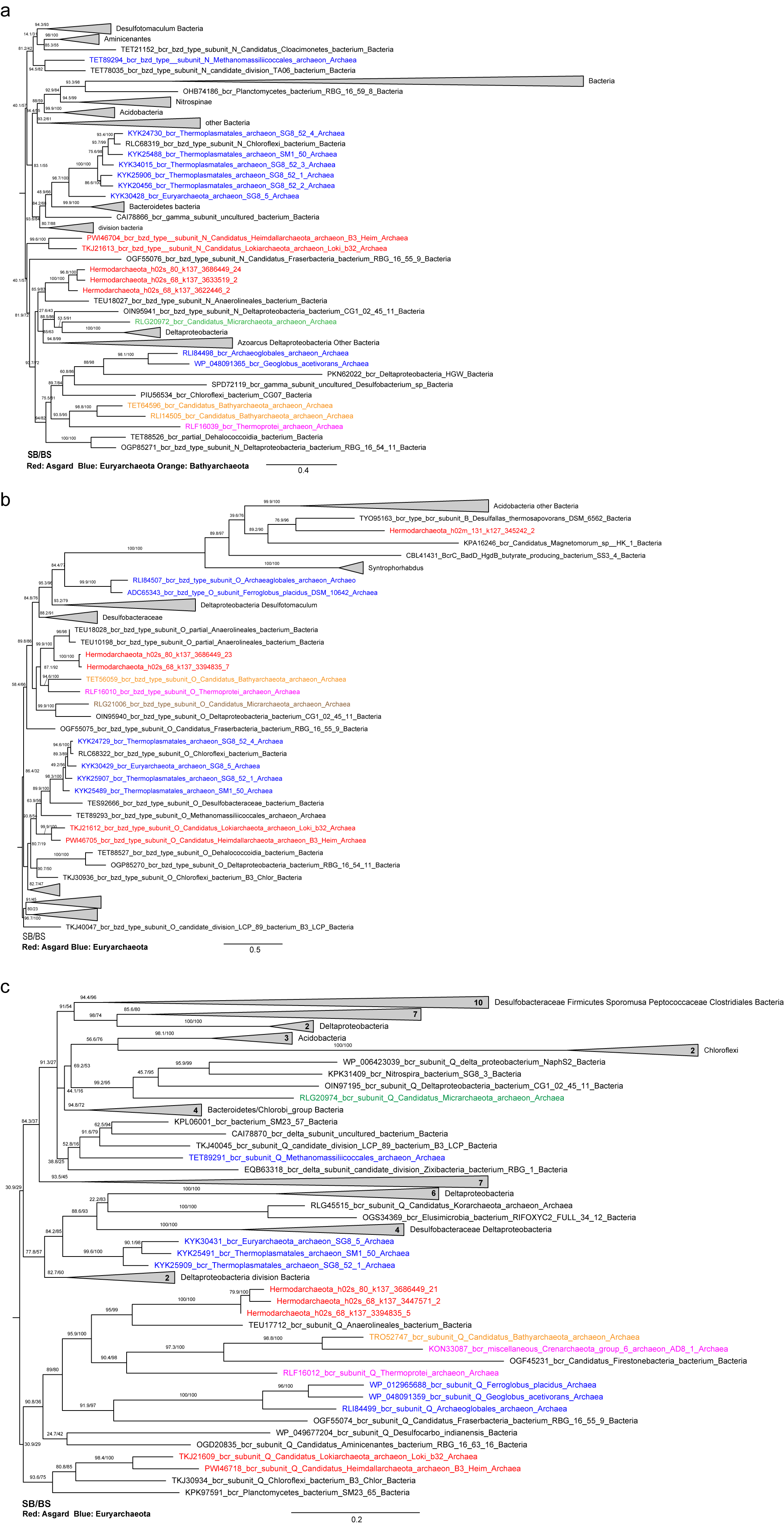

**Supplementary Fig. 13** Maximum-likelihood tree of benzoyl-CoA reductase (bcr) identified in Hermodarchaeota genomes and homologues from nr database reconstructed using IQtree with LG+G4 model. Asgard archaeal benzoyl-CoA reductases were red-coded. **a** Benzoyl-CoA reductase subunit N; **b** Benzoyl-CoA reductase subunit O; **c** Benzoyl-CoA reductase subunit Q. The SH-like approximate likelihood ratio test (SB) and ultrafast bootstrap (BS) support values were shown.

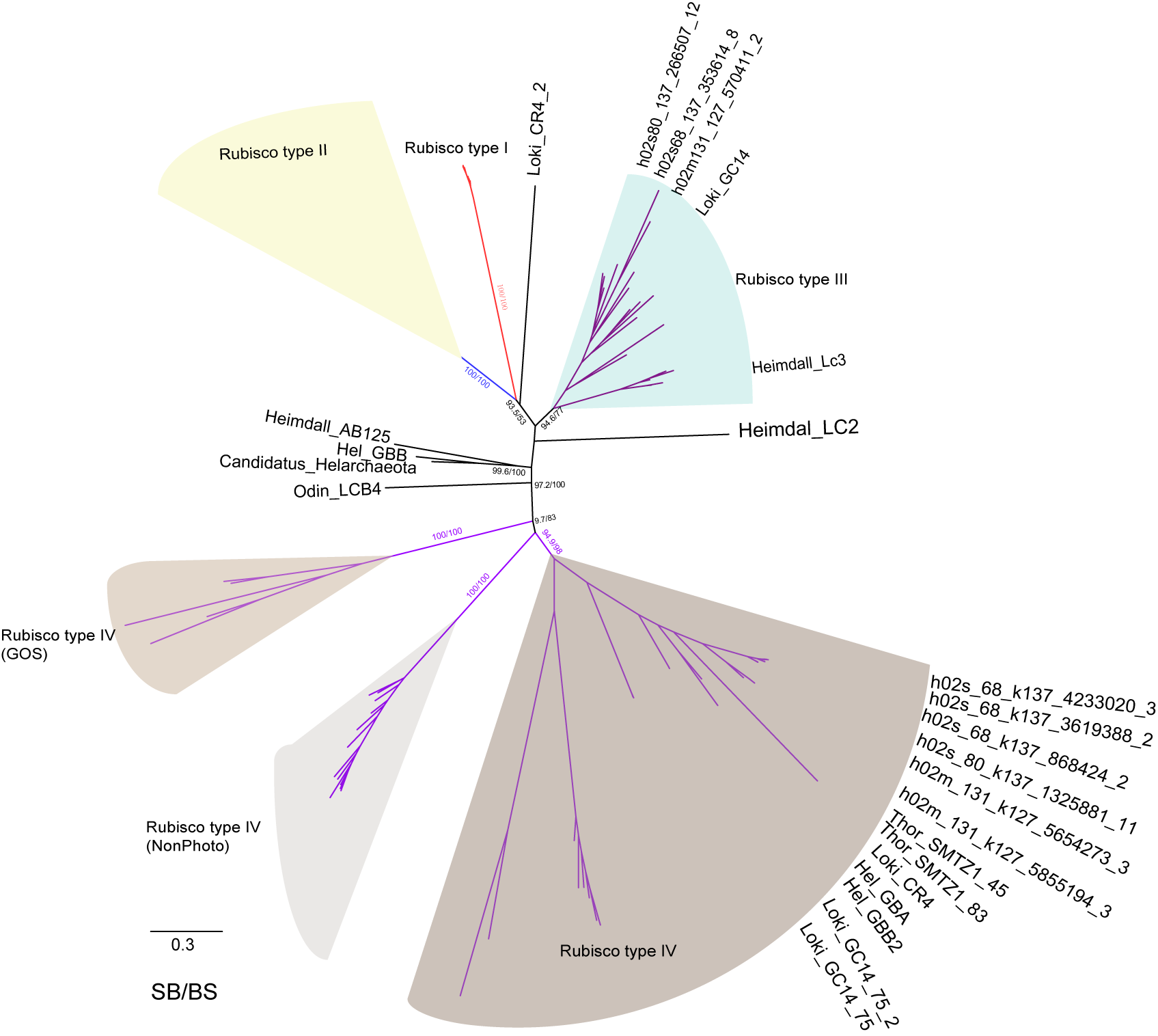

**Supplementary Fig. 14** Maximum-likelihood tree of 1,5-bisphosphate carboxylase/oxygenase (Rubisco) identified in Hermodarchaeota genomes. Other Rubisco sequences refer to previous studies [5, 15]. Maximum-likelihood tree was reconstructed using IQtree with LG+I+G4 model. This tree indicates the four types of Rubisco proteins. The SH-like approximate likelihood ratio test (SB) and ultrafast bootstrap (BS) support values were shown.
